# Novel Driver Mutations in GCB Lymphoma Patients That Affect Transcription Factors Binding

**DOI:** 10.1101/2025.03.15.643417

**Authors:** Ofek Shami-Schnitzer, Tamir Tuller

## Abstract

Mutations in the DNA can affect cancer development and progression not only by changing the amino acid chain but also by affecting different regulatory elements such as transcription factors. This study introduces a novel pipeline to identify “mutation blocks” - small genomic areas with high mutation rates that potentially influence transcription factors binding. By analyzing GCB lymphoma patient data, mutations blocks were identified that correlated with gene expression changes and were linked to transcription factor activity. These mutation blocks suggest a selection for mutations that alter gene regulation, contributing to lymphoma development. The analysis identified 56 mutation blocks in germinal center B-cell like diffuse large B-cell lymphoma (GCB DLBCL) patients’ genomes, affecting genes such as BCL2, MYC, SGK1, and PIM1, and linked to transcription factors including MSC, TCFL5, HOXB7, FOXP3, and ZBTB6. A machine learning model that used gene ontology data suggested further potential transcription factors-gene pairs influencing cancer. These findings highlight the role of synonymous and silent mutations in altering transcription factors binding and gene expression, offering insights into the mechanisms of GCB lymphoma.

## 1 Introduction

Mutations in DNA are considered a central driver of neoplasm development and, consequently, cancer [1]. While some mutations are found in the coding region of the gene and may alter the protein by changing its amino acid sequence, other mutations, such as silent mutations, do not affect protein structure.

Regardless of its effect on the amino acid sequence, a mutation might affect gene regulatory mechanisms and contribute to cancer development[2−4]. Previous studies have shown mutations that disrupt exon splicing sites[5, 6]. Other studies have identified synonymous mutations in the coding region that replace nucleotides without altering the amino acid sequence but still affect regulatory processes[7−9].

Transcription factors (TFs) are proteins that regulate gene transcription by binding to specific DNA sequences. Depending on their function, transcription factors may either activate transcription (activators) or suppress it (repressors). This regulatory mechanism plays a significant role in cancer development[10, 11]. Thus, mutations in a gene may have little to no effect on the resulting protein but can significantly impact transcription factor binding sites, thereby altering gene regulation[12−14].

The BCL2 family comprises a group of genes responsible, for regulating apoptosis— the intrinsic cell death mechanism. One of its key members is the BCL2 (B-Cell lymphoma 2) gene, which functions as a pro-survival gene by inhibiting the release of cytochrome C. This, in turn, prevents the activation of caspase-9, thereby suppressing apoptosis. Due to its major role as a pro-survival gene, malfunction of BCL2 may interfere with the apoptosis system [15, 16] and overexpression of BCL2 is known to be connected to large diffuse B-cell lymphoma (DLBCL), the most frequent B cell Non-Hodgkin’s lymphoma. The translocation BCL2(14;18)(q32;q21) is a common trait among DLBCL patients. This translocation found to be the cause of 20-30% of over-expression of BCL2 among DLBCL patients [17, 18]. Moreover, the translocation BCL2(14;18)(q32;q21) is known to increase the creation of mutations on the BCL2 gene (hypermutation), synonymous mutations included[17]. Germinal Center B Cell-Like (GCB) is a sub-group of DLBCL, distinguished by gene expression profiling[19].

A previous study showed that among GCB lymphoma patients, two recurrent synonymous mutations in the coding region of the BCL2 gene may affect MSC repressor binding[8]. In this study, we did not focus on specific mutations but instead identified small genomic regions with high mutation density, which we termed mutation blocks. Since transcription factors bind to short DNA sequences, we hypothesize that different mutations within the same region may similarly impact cancer development by altering transcription factor binding sites. Therefore, we analyzed large-scale genomic data, including data from GCB lymphoma patients, to identify mutation blocks that may alter transcription factor binding potential.

## 2. Results

### 2.1 A pipeline for finding mutations affecting TF binding sites

A multistage data pipeline was developed to identify genomic regions and associated transcription factors that together exhibit all of the following features:

- These regions have a high mutation rate in cancer patients, suggesting positive selection for mutations in these sites
- Mutations in these regions correlate with changes in gene expression within the same genomic region
- A transcription factor-related mechanism may explain these changes We refer to these genomic regions as mutations blocks.

The pipeline begins with the **selection identification** stage, where mutations blocks (genomic regions with a high mutation rate in cancer patients) are first identified. Next, in the **RNA validation** stage, we identify mutation blocks where mutations correlate with changes in gene expression within the same region. Finally, we select mutation blocks that affect gene expression and proceed to the **transcription factor analysis** stage. This stage consists of three tests. The first test, **tissue validation**, ensures that the transcription factor is active in the same tissue where the cancer occurs. The second test, **mechanism validation**, verifies that the transcription factor’s role (activator or repressor) aligns with the observed gene expression changes from the RNA validation stage. The final test, **binding change verification**, calculates the predicted change in the transcription factor’s binding affinity (either stronger or weaker binding to the genome). Using this predicted binding change, we assess whether the observed gene expression effects are biologically plausible.

The final output of this pipeline consists of trios, each containing three components:

- **Mutation blocks** with an increased mutation rate in a specific cancer type.
- **Genes** where these mutation blocks are located and show altered expression.
- **Transcription factors influenced** by these mutations, which are active in the relevant tissue and have a plausible regulatory mechanism.

A visualization of the pipeline is provided in Fig. 1. All the details appear in the methods section.

**Fig 1.**
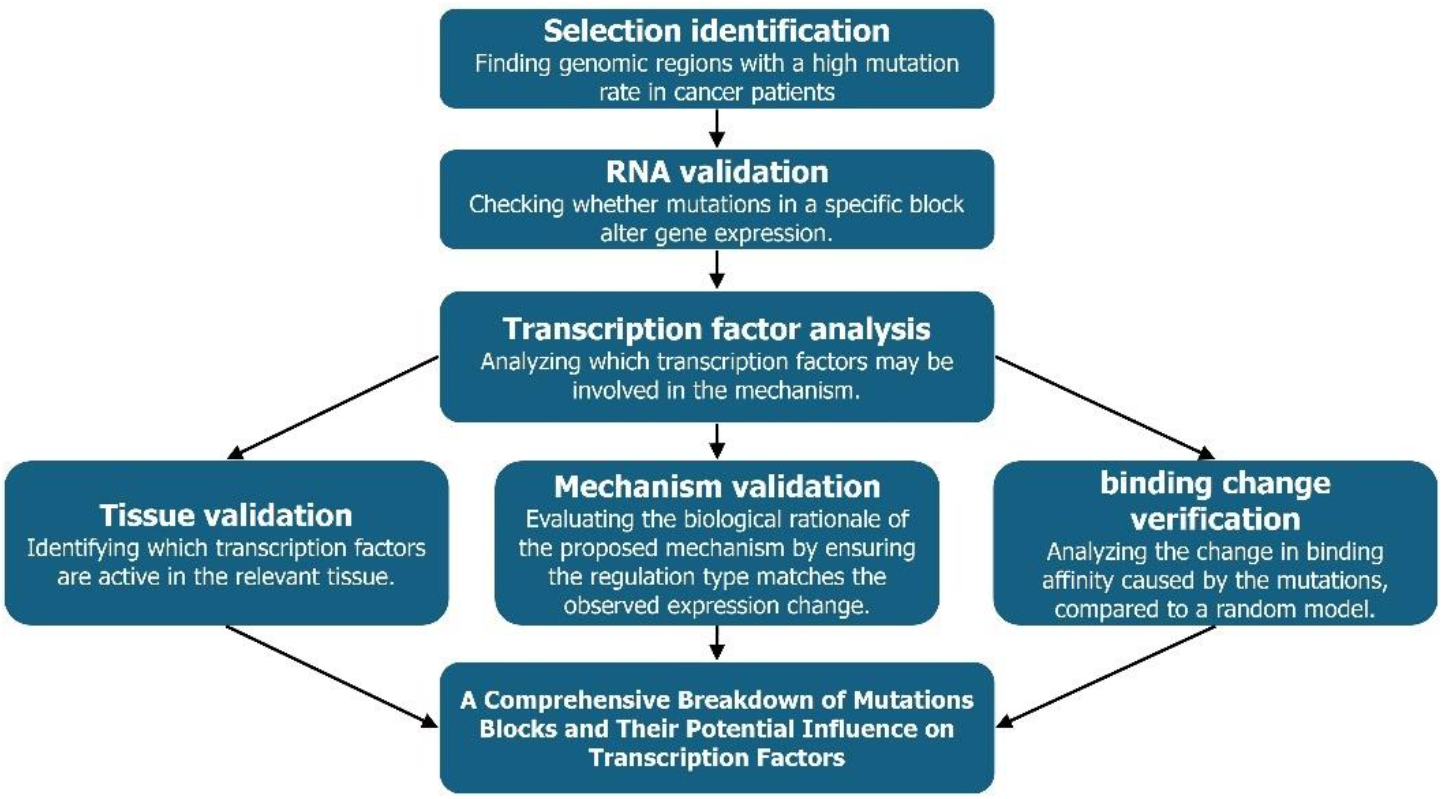
a visualization of the pipeline for finding mutations effecting TF binding sites. Each block in the visualization represents the different stages of the pipeline.

### 2.2. Systems biology study of mutations blocks

#### Our analysis is based on two assumptions

1. **Mutations in the binding sites of various transcription factors may play a key mechanism in the development of GCB Lymphoma** – a selection toward numerous binding sites mutations blocks in different genes were found in GCB lymphoma patients. Mutations within these blocks are correlated with changes in the expression of the relevant genes. In addition, mutations in these mutations blocks Result in significant changes in the predicted binding affinity of transcription factors to the gene. Given the extensive number of mutations blocks identified, we hypothesize that this mechanism (mutations in the binding sites of transcription factors) is a key driver in GCB lymphoma.
2. **Certain genes and transcription factors may play a crucial role in the development of GCB lymphoma** – in some genes (BCL2, MYC, and SGK1) Multiple mutations blocks in potential binding sites have been identified. For some transcription factors (FOXP3, HOXB7, MSC, TCFL5 and ZBTB6) multiple sites of mutations blocks with binding potential for these transcription factors were found. The recurrence of these genes and transcription factors with mutation blocks exhibiting a high mutation rate in GCB patients suggests their key role in GCB lymphoma development.

The pipeline output includes 56 distinct mutations blocks identified in GCB patients. These mutations blocks are located in oncogenes with a high concentration of mutations. Patients with mutations in these blocks showed significant changes in the expression of the corresponding gene. In addition, Additionally, a plausible transcription factor-related mechanism was identified for each of these mutations blocks. Therefore, we hypothesize that these mutation blocks alter the binding affinity of specific transcription factors, thereby affecting gene expression.

Certain genes and transcription factors were identified multiple times in the results. suggesting their broader role in GCB DLBCL Their expression was affected in several distinct locations across the genome. A heatmap of the 56 mutations blocks, the genes in which they were found and the transcription factors that they might affect is on Fig. 2. Note that each mutations block affects a single gene but may influence the binding of multiple transcription factors. Several mutation blocks were identified in the following **genes**:

**BCL2** − out of 56 mutations blocks identified in the results, 14 are located along the BCL2 gene. BCL2 is a regulator that prevents apoptosis, the cellular death mechanism[15]. Mutation in all 14 mutations blocks are corelated with upregulation of BCL2 and thereby preventing cell death. Additionally, BCL2 is known for its association with DLBCL[18].

**MYC** − Five mutations blocks were identified along the MYC gene. MYC is an oncogene that influences various cellular processes, including apoptosis, DNA damage response, and the cell cycle[20]. All of the mutations blocks found in MYC lead to overexpression of the gene, which is also associated with DLBCL[21, 22].

**SGK1** − Another five mutation blocks were identified on the SGK1 gene, leading to its overexpression. SGK1 is a protein kinase whose malfunction is associated with various types of cancer[23]. Additionally, SGK1 is known to generate hyper-stable proteins in DLBCL patients, which affect the progression of DLBCL[24].

**PIM1** − Three of the 56 mutation blocks were identified in the PIM1 gene. PIM1 is a serine/threonine kinase that is overexpressed in many human cancers[25] including DLBCL[26]. All three mutations blocks found in PIM1 lead to upregulation of the gene.

**Fig 2.**
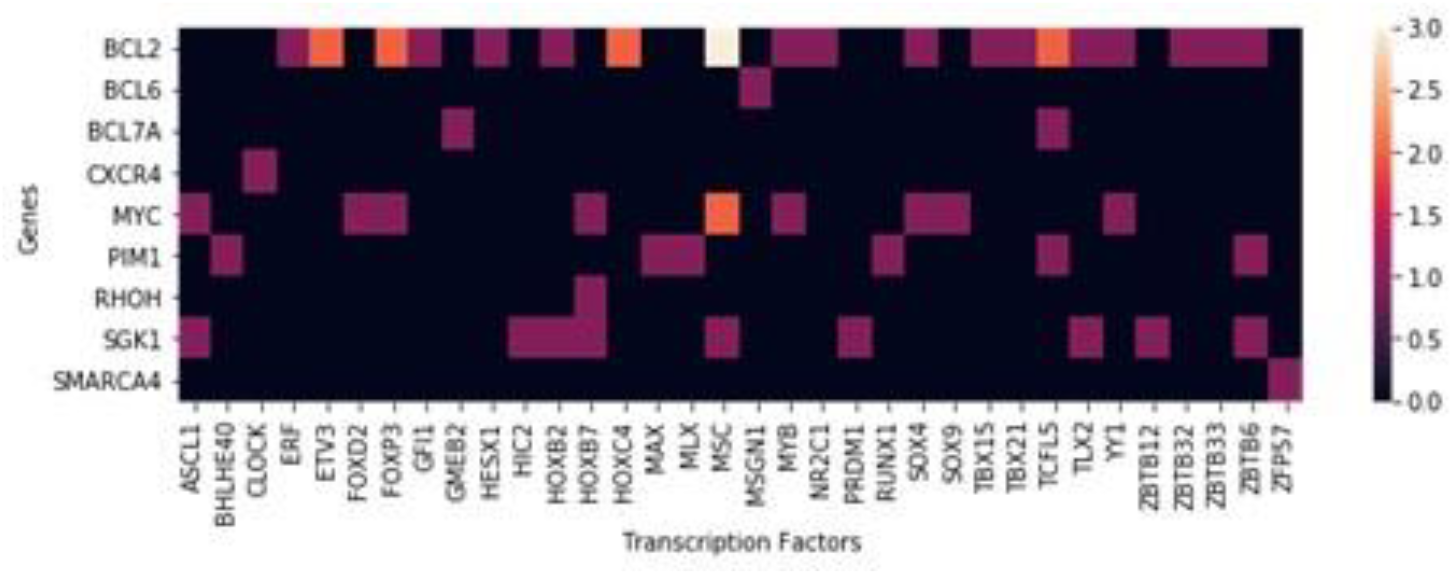
A heatmap representing the possible connection between genes and transcription factors across GCB lymphoma patients. The color intensity indicates the frequency (number of times) where each transcription factor was found to be potentially connected to a mutations block in a specific gene. For example, two mutation blocks in the BCL2 gene were found to be potentially connected to TCFL5, which was also identified as potentially connected to mutation blocks in the BCL7A and PIM1 genes.

The following **transcription factors** were identified as potential contributors to multiple mutation blocks:

**MSC** (also known as ABF1) − Six of the 56 mutation blocks were found to potentially affect the binding of the MSC repressor. MSC is a transcription factor known to be active in lymphocytes[27] and has previously been associated with lymphoma[28]. These six mutation blocks may lead to overexpression of BCL2 (three blocks), MYC (two blocks), and SGK1.

**TCFL5** − four mutations blocks were found to potentially affect TCFL5 binding. The TCFL5 repressor plays an important role in spermatogenesis, the immune system, and cancer[29]. In all four mutation blocks, the binding affinity of TCFL5 is reduced, leading to overexpression of BCL2 (two blocks), PIM1, and BCL7A.

**HOXB7** − three mutations blocks out of 56 found to potentially affect the binding of HOXB7 activator. HOXB7 regulates a variety of genes and several oncogenic pathways, and it is overexpressed in cancer[30] and has also been correlated with DLBCL[31]. Mutation in all three mutation blocks are corelated with increase in the binding of HOXB7, leading to the overexpression of the genes RHOH, MYC, and SGK1.

**FOXP3** − three out of the 56 mutations blocks found to potentially affect the binding of the repressor FOXP3. FOXP3 induces immunosuppressive functions in cells[32] and is highly expressed in the tissues of DLBCL patients[33]. Two of these mutation blocks are located along the BCL2 gene, and one is on MYC, all of them reduce the binding affinity of the FOXP3 repressor, leading to overexpression of the BCL2 and MYC genes.

**ZBTB6** − three mutations blocks found to potentially affect the binding of ZBTB6 repressor. ZBTB6 is a member of the BTB-ZF transcription factor family and has been connected to B-cell development[34]. Mutations in all three mutations blocks are corelated with overexpression of different genes: BCL2, PIM1 and SGK1.

#### BCL2 example of the pipeline results

As an example of the pipeline results, a single mutation block analysis will be presented. Additional analyses of five mutation blocks related to MSC can be found in the supplementary data section 2, along with a csv table summarizing all 56 mutations blocks.

Mutations block 63319682 − MSC binding affecting BCL2:

This mutations block includes 14 different mutations in the 5’ prime UTR that were found 32 times (among 241 GCB DLBCL patients). The block is located on chromosome 18: 63319682-63319692, where the BCL2 gene is found. A detailed representation of the identified mutations is provided in Fig. 3.

**Fig 3.**
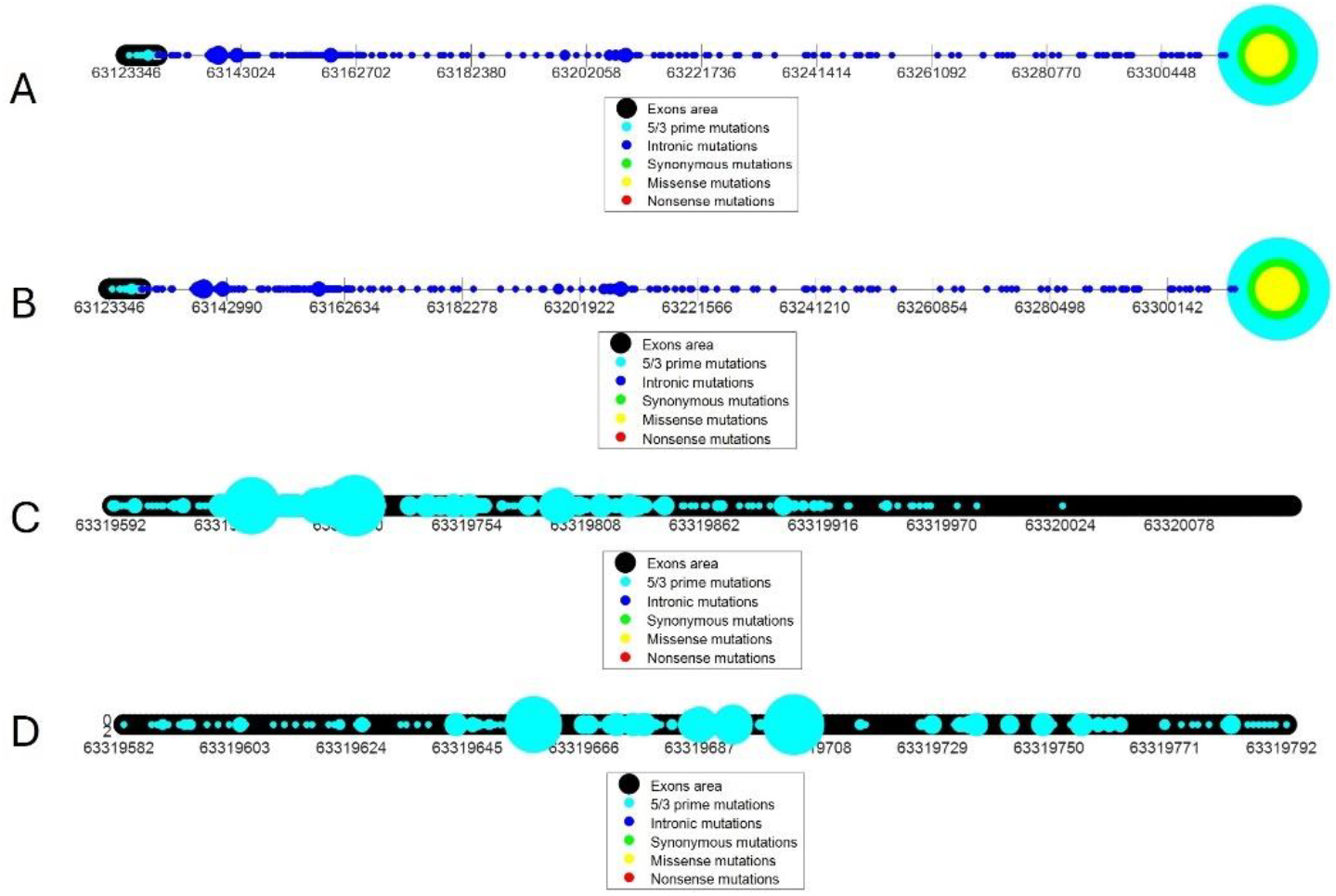
This figure illustrates the distribution of mutations in GCB DLBCL patients along the BCL2 gene with circle sizes representing mutation frequency. The figure is divided into four sections: (A) mutations across the entire gene, (B) mutations from the gene’s start to the mutation block at positions 63319682−63319692 (which is adjacent to the end of the gene), (C) mutations from the mutation block at positions 63319682−63319692 to the gene’s end, and (D) mutations within ±100 nucleotides of the mutation block. As shown, the 63319682−63319692 mutation block contains several mutations in the 5’ prime UTR, some of which are highly recurrent.

To validate the effect of this mutation on gene expression, BCL2 expression levels were analyzed. Patients with the mutations exhibited significantly higher BCL2 expression compared to randomized groups of GCB lymphoma patients (P-value = 0.0013; see Fig. 4) This overexpression of BCL2 may contribute to cancer progression due to its anti-apoptotic function.

**Fig 4.**
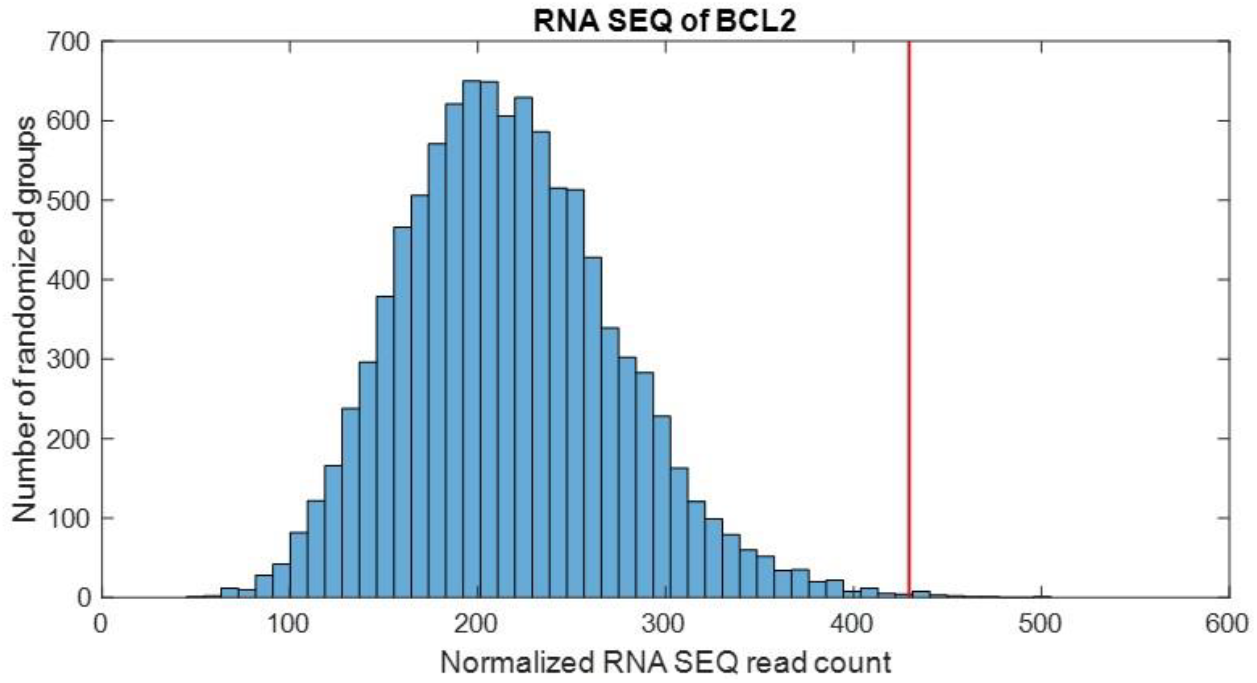
Using RNA sequencing data, 10,000 random patient groups were selected, and their mean expression levels were calculated. The red line represents the mean expression of patients with mutations in the 63319682-63319692 block of the BCL2 gene.

A search for relevant transcription factors that could explain the observed change in expression was conducted. Only transcription factors expressed in EBV-transformed lymphocytes were considered and analyzed for changes in their binding ability to the relevant genomic region. One transcription factor was identified as a potential cause of the observed BCL2 overexpression: MSC. The mutations in the identified mutation block interfere with the binding of the MSC repressor to BCL2, as shown in Fig. 5, potentially leading to BCL2 overexpression.

**Fig 5.**
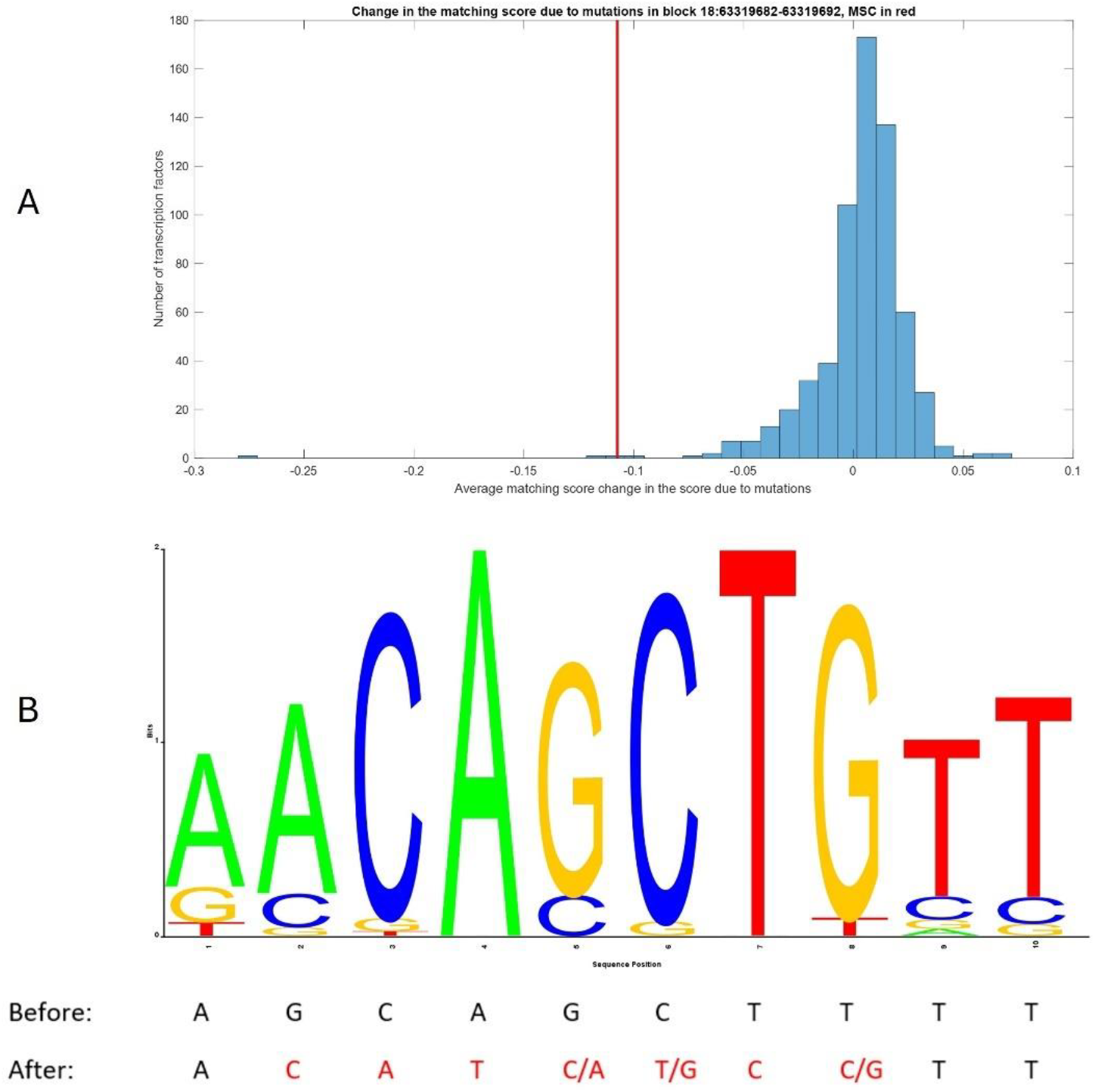
illustrates the effect of mutations on the ability of transcription factors to bind to the BCL2 gene. In A, a histogram representing the impact of mutations within the identified block on the binding ability of various transcription factors. As shown, most transcription factors are minimally affected by these mutations, while a few exhibits significant binding changes. MSC is highlighted with a red line. In B, the sequence LOGO of the MSC binding site, based on JASPAR’s Position-Specific Scoring Matrices (PSSMs). Below, the nucleotide sequence of the relevant site (18: 63319680-63319689, reverse strand) is shown with the mutation (after mutation) and without the mutation (before mutation). Alternative nucleotides found in patients are marked in red.

### 2.3 Gene ontology is associated with the reported TF-gene induced by cancer mutations

We suspected that additional TF-gene pairs influencing cancer may have been missed due to data limitations. To address this, we applied machine learning techniques to identify potential TF-gene pairs that were not detected in our initial analysis. To achieve this, we utilized two distinct datasets, both separately and in an integrated manner:

1. Gene Ontology (GO)[35, 36] as the connection between transcription factor and gene can be examined by assessing their functional relationships.
2. Protein-Protein Interactions (PPI)[37] as the connection between transcription factor and gene can be examined by assessing their functional relationships based on the relationships between different proteins

For each method (GO and PPI), a graph was constructed: for PPI, a graph of genes, and for GO, a graph of terms and genes. For each node in these graphs, a vector representation was generated using the Node2Vec algorithm[38]. Each possible transcription factor-gene pair was then represented by a joint vector, created by merging the respective vectors of the transcription factor and the gene. The previously identified cancer-related mutation blocks in genes were used as positive data, while pseudo-negative data was generated. Finally, a LightGBM[39] supervised learning model was trained on this data and evaluated for accuracy. To verify the significance of the results, another GO graph was created, this time with randomized associations between gene/term and their respected vectors.

Using only GO data, the model was able to predict gene-transcription factor pairs with a modest accuracy (all results above 0.5, best accuracy result is 0.589). In contrast, the randomised GO version and using only PPI data did not yield any significant results. When the two datasets (GO and PPI) were merged, the results were similar to those obtained using only GO data (see full results in supplementary data table 1). These findings suggest that the functional relationship between the gene and transcription factor, as represented by the GO data, provides some predictive value for their potential connection to cancer. However, their ability to physically bind, as represented by the PPI data, did not contribute significantly to the prediction.

## 3 Discussion

The findings of this study highlight the critical role that synonymous and silent mutations, particularly within transcription factor binding sites, play in the development of GCB lymphoma. Traditionally, the focus of cancer genomics has been on mutations in coding regions that directly alter protein function. However, this study expands the perspective by demonstrating that synonymous mutations or those in synonymous and silent regions can have a profound impact by modifying gene regulation through changes in transcription factors binding affinity.

The identification of 56 mutation blocks in GCB DLBCL patients highlights the significance of clustered mutations in specific regions of the genome. These mutations, correlated with changes in gene expression, suggest a selective pressure in cancer development towards disrupting expression using regulatory mechanisms. Notably, the study points to genes such as BCL2, MYC, SGK1, and PIM1—all known to be involved in oncogenesis. The change in expression of these genes, likely driven by disrupted transcription factors binding, aligns with their established roles in promoting cell survival, proliferation, and tumor progression.

An important contribution of this work is its focus on the transcription factors MSC, TCFL5, HOXB7, FOXP3, and ZBTB6. These transcription factors are already known to play a role in cancer or lymphocytes, but the discovery of multiple mutation blocks affecting their binding sites suggests that these transcription factors might have even broader implications in GCB DLBCL pathogenesis. For example, the recurrent mutation blocks found in BCL2 associated with MSC binding, support a model in which disrupted transcription factors regulation contributes to their overexpression and thus, to lymphoma development. These TFs may serve as potential therapeutic targets or biomarkers, as their activity seems to be linked to key oncogenic processes in lymphoma.

Furthermore, the use of machine learning to predict additional transcription factors-gene pairs based on gene ontology (GO) data represents a novel approach that can expand the scope of the study beyond the available dataset. The modest predictive success of gene ontology-based models indicates that while functional relationships between genes and transcription factors can inform their cancer relevance, further refinement in the model—perhaps integrating more nuanced datasets or additional biological features—may be needed to improve predictions. The lack of signal in the protein-protein interaction (PPI) dataset is intriguing and suggests that the functional relationship between gene and transcription factors as reflected by interactions between proteins, may not be as critical in this context as their functional associations. This may be related to the fact that transcription factors mainly interact with DNA molecules and not proteins and/or to noise and bias in PPI networks.

One of the most significant implications of these findings is the potential for clinical translation. Identifying mutation blocks that disrupt transcription factors binding opens new avenues for precision oncology. Drugs targeting specific transcription factors or the pathways they regulate could be developed or repurposed to counteract the deregulation caused by these mutations. Moreover, the mutation blocks identified in this study could serve as biomarkers for GCB DLBCL prognosis or treatment response, allowing for more personalized therapeutic strategies.

However, there are some limitations to consider. The study’s reliance on genomic data from a specific cancer type, GCB lymphoma, means that the generalizability of the findings to other cancers remains uncertain. Additionally, while correlations between mutation blocks, transcription factors binding, and gene expression changes are compelling, further experimental validation is needed to confirm the causal role of these mutations in altering transcription factors binding and driving cancer progression. Functional assays could be employed to directly measure transcription factors binding in these regions.

In conclusion, this study provides compelling evidence that mutations, including synonymous and silent mutations which affect transcription factors binding sites, are key drivers in GCB lymphoma. The pipeline developed here offers a robust framework for identifying similar mutation blocks in other cancers, paving the way for deeper insights into cancer regulatory mechanisms. Future work should focus on validating these findings experimentally and exploring the therapeutic potential of targeting the transcription factors implicated in these mutation blocks.

## 4. Methods

A pipeline was developed to detect driver mutations in cancer that affect transcription factor binding. This pipeline processes patient mutation data from a specific cancer type using a step-based method to identify relevant mutations. First, we search for areas in the genome with a high mutation rate, particularly within cancer-related genes. A high mutation rate suggests selective pressure in cancer development. Next, we validate, using RNA sequencing data, that mutations in these regions—referred to as mutation blocks—are correlated with changes in gene expression, as expected for transcription factor-related mutations. We then examine which transcription factors might be impacted by these mutation blocks, based on their binding ability and the effect of mutations on this binding. Lastly, we apply three validation methods to ensure the pipeline outputs relevant mutations: confirming that the transcription factor is active in the tissue relevant to the cancer type, verifying the biological logic of the proposed mechanism, and validating the significance of the binding affinity change. A detailed explanation of each step follows. A visualization of the method is shown in Fig. 1.

## 4.1 Selection identification

Data from patients with malignant lymphoma, specifically those with germinal center B-cell derived lymphomas (GCB), were obtained from the ICGC-MMML-Seq (Molecular Mechanisms in Malignant Lymphoma by Sequencing) Consortium[40]. This dataset includes simple somatic mutations (SSMs) from 241 patients and sequence-based gene expression data for 105 patients. Initially, we performed a preprocessing step to enhance our SSM data. During this step, we converted the mutation locations in the genome from GRCh37 to GRCh38 using the UCSC Genome Lift Tool[41].

To identify the mutations blocks − areas in the genome with high mutations rate we have used the following process. First, for each chromosome all mutations of GCB patients were acquired. Then the initial blocks were created using a sliding window in the size of 10 nucleotides. Afterwards, high mutated blocks were chosen by taking only 5% of blocks with the highest number of mutations (out of blocks that have one or more mutations). Lastly all high mutated blocks that were overlapping blocks were merged into a bigger block, and these are the final blocks that resemble areas with high mutations rate in GCB patients.

### 4.2 RNA validation

Using the highly mutated blocks and cancer related genes database (COSMIC[42]) we identified cancer related genes that are found in the blocks’ locations. Then, going over each of these genes and using RNA-seq data from the ICGC of GCB patients a data set of the expression of certain gene per patient is created. We then calculated the mean expression of the gene for patients with a mutation in the block area, is found. In parallel, a random model of expression was created by randomly selecting groups of patients (same size as the number of patients with mutations in the relevant block) and calculating their mean expression. We repeated this process 10,000 times to generate a random model that was created and the patients with mutations in the block mean expression were measured against it to calculate p-value.

### 4.3 Transcription factor analysis

This step is composed of three different sub-steps: the binding grade change verification, tissue validation and mechanism understanding. Together, these sub-steps provide a comprehensive approach that is greater than the sum of its parts. These three results help identify transcription factors whose binding is significantly impacted by the mutations in the mutations block (binding grade change verification), that are active in the relevant tissue (tissue validation) and that their mechanism of action is consistent with the effect seen in the previous expression test.

### 4.4 Binding change verification

In this step the effect of the mutations on the binding ability of different transcription factors is tested. Using JASPAR’s[43] PSSM binding data, we acquired the binding score (which represents the binding probability) of each human transcription factor (746 in total) to the mutations block area without the mutations and compared it to the mean score of the mutated block (by applying each mutation independently and normalizing by the number of mutations found in patients). Afterwards we considered only transcription factors that were highly affected by the mutations in the mutations block, by considering only transcription factors that their change in the binding score has a z score higher than three or lower than minus three (using all other transcription factors as a background model).

### 4.5 Tissue validation

This step is crucial in order to validate that the transcription factor found is indeed relevant to this cancer type. To this end, GTEx database, which provides expression data across various tissues[44] was used to validate whether the highly influenced transcription factors from the binding grade verification are indeed present in the relevant tissue − lymphocytes, compared their expression levels to other tissues.

### 4.6 Mechanism validation

Finally, we assessed if whether the proposed mechanism, in which mutations in transcription factor binding sites influence gene expression, is biologically valid. Therefore, using gene ontology[35, 36, 45] the transcription factor type (either it is an activator or a repressor) was found. Then, we took each trio of transcription factor type, binding probability change, and gene expression change and checked if the trio form a biologically plausible mechanism. For example, if the binding of a repressor was harm and an increase in the gene expression was found − the entire suggested mechanism makes a biologically reasonable mechanism.

### 4.7 System biology analysis

Finally, a macro-level analysis of the results was made as a means to discover the broad effects of these mechanisms. In this analysis the results from all chromosomes were aggregated and grouped according to transcription factors and influenced genes. This approach revealed transcription factors and genes that were recurrently affected across the dataset.

### 4.8 Machine Learning

To expand our results and identify more possible duos of transcription factors and gens that are influenced by our suggested mechanism, a machine learning approach was employed. We tried four different approaches by constructing two distinct graphs:

Gene Ontology (GO) − We constructed a graph of all GO terms[35, 36, 45] and then added to this graph all genes found in GO by incorporated genes associated with each term by adding them as nodes. Each gene node was then connected to the relevant GO terms. Additionally, for each cancer-related gene from COSMIC[42], if it was not already represented in the graph, we manually added it (without linking it to any GO term).

Protein-Protein Interactions (PPI) - Similar to the GO graph, we constructed a graph of all proteins and their interactions[37]. Then, we added cancer-related genes from COSMIC that were not already represented as new nodes.

Afterwards we performed a grid search with different parameters (vector length of 8, 16 and 32; walk length of 4, 8 and 16; window size of 4,8 and 16). For each group of parameters in the grid search, the following steps were taken. First we created two node2vec[38] models, one for each graph (GO and PPI). Then we represented each gene by his vector representation of its place in the graph using four different approaches − GO, PPI, merged approach (by concatenating the two previous vectors) and randomized GO in which after generating vectors for each node, we shuffled the node− vector assignments. Lastly we trained a LGBM model[39]. For positive examples we took all gene-transcription factors duos found in our study. For pseudo negative examples we took random pairs of gene-transcription factors (taking into consideration that there are millions of possible duos and only a few hundred were found in our research). To the end of measuring our results we performed 10-fold cross validation test, ensuring no leakage between the training and test sets in each fold. Specifically, a gene would appear in either the training or the test set, but not both, and the same rule applied to each transcription factor.

## Acknowledgments

The study was partially supported by the koret-berkeley-tau initiative.

## References

1. Futreal, P.A., Coin, L., Marshall, M., Down, T., Hubbard, T., Wooster, R., Rahman, N., Stratton, M.R.: A census of human cancer genes. Nat. Rev. Cancer. 4, 177 (2004).

2. Diederichs, S., Bartsch, L., Berkmann, J.C., Fröse, K., Heitmann, J., Hoppe, C., Iggena, D., Jazmati, D., Karschnia, P., Linsenmeier, M., Maulhardt, T., Möhrmann, L., Morstein, J., Paffenholz, S.V., Röpenack, P., Rückert, T., Sandig, L., Schell, M., Steinmann, A., Voss, G., Wasmuth, J., Weinberger, M.E., Wullenkord, R.: The dark matter of the cancer genome: aberrations in regulatory elements, untranslated regions, splice sites, non-coding RNA and synonymous mutations. EMBO Mol. Med. 8, 442 (2016). 10.15252/emmm.201506055.

3. Gutman, T., Goren, G., Efroni, O., Tuller, T.: Estimating the predictive power of silent mutations on cancer classification and prognosis. Npj Genomic Med. 6, 67 (2021). 10.1038/s41525-021-00229-1.

4. Zhang, X., Meyerson, M.: Illuminating the noncoding genome in cancer. Nat. Cancer. 1, 864−872 (2020). 10.1038/s43018-020-00114-3.

5. Cartegni, L., Chew, S.L., Krainer, A.R.: Listening to silence and understanding nonsense: exonic mutations that affect splicing. Nat. Rev. Genet. 3, 285−298 (2002). 10.1038/nrg775.

6. Lynn, N., Tuller, T.: Detecting and understanding meaningful cancerous mutations based on computational models of mRNA splicing. Npj Syst. Biol. Appl. 10, 25 (2024). 10.1038/s41540-024-00351-7.

7. Gartner, J.J., Parker, S.C.J., Prickett, T.D., Dutton-Regester, K., Stitzel, M.L., Lin, J.C., Davis, S., Simhadri, V.L., Jha, S., Katagiri, N., Gotea, V., Teer, J.K., Wei, X., Morken, M.A., Bhanot, U.K., Chen, G., Elnitski, L.L., Davies, M.A., Gershenwald, J.E., Carter, H., Karchin, R., Robinson, W., Robinson, S., Rosenberg, S.A., Collins, F.S., Parmigiani, G., Komar, A.A., Kimchi-Sarfaty, C., Hayward, N.K., Margulies, E.H., Samuels, Y.: Whole-genome sequencing identifies a recurrent functional synonymous mutation in melanoma. Proc. Natl. Acad. Sci. 110, 13481 (2013). 10.1073/pnas.1304227110.

8. Shami-Schnitzer, O., Zafir, Z., Tuller, T.: Novel Driver Synonymous Mutations in the Coding Regions of GCB Lymphoma Patients Improve the Transcription Levels of BCL2. In: Bebis, G., Alekseyev, M., Cho, H., Gevertz, J., and Rodriguez Martinez, M. (eds.) Mathematical and Computational Oncology. pp. 108−118. Springer International Publishing, Cham (2020).

9. Gutman, T., Tuller, T.: Computational Analysis of MDR1 Variants Predicts Effect on Cancer Cells via their Effect on mRNA Folding. PLOS Comput. Biol. 20, e1012685 (2024). 10.1371/journal.pcbi.1012685.

10. Nebert, D.W.: Transcription factors and cancer: an overview. Toxicology. 181−182, 131− 141 (2002). 10.1016/S0300-483X(02)00269-X.

11. Bushweller, J.H.: Targeting transcription factors in cancer — from undruggable to reality. Nat. Rev. Cancer. 19, 611−624 (2019). 10.1038/s41568-019-0196-7.

12. Morova, T., McNeill, D.R., Lallous, N., Gönen, M., Dalal, K., Wilson, D.M., Gürsoy, A., Keskin, Ö., Lack, N.A.: Androgen receptor-binding sites are highly mutated in prostate cancer. Nat. Commun. 11, 832 (2020). 10.1038/s41467-020-14644-y.

13. Katainen, R., Dave, K., Pitkänen, E., Palin, K., Kivioja, T., Välimäki, N., Gylfe, A.E., Ristolainen, H., Hänninen, U.A., Cajuso, T., Kondelin, J., Tanskanen, T., Mecklin, J.-P., Järvinen, H., Renkonen-Sinisalo, L., Lepistö, A., Kaasinen, E., Kilpivaara, O., Tuupanen, S., Enge, M., Taipale, J., Aaltonen, L.A.: CTCF/cohesin-binding sites are frequently mutated in cancer. Nat. Genet. 47, 818−821 (2015). 10.1038/ng.3335.

14. Kaiser, V.B., Taylor, M.S., Semple, C.A.: Mutational Biases Drive Elevated Rates of Substitution at Regulatory Sites across Cancer Types. PLOS Genet. 12, e1006207 (2016). 10.1371/journal.pgen.1006207.

15. Cory, S., Adams, J.M.: The Bcl2 family: regulators of the cellular life-or-death switch. Nat. Rev. Cancer. 2, 647 (2002).

16. Vaux, D.L., Cory, S., Adams, J.M.: Bcl-2 gene promotes haemopoietic cell survival and cooperates with c-myc to immortalize pre-B cells. Nature. 335, 440 (1988).

17. Lohr, J.G., Stojanov, P., Lawrence, M.S., Auclair, D., Chapuy, B., Sougnez, C., Cruz-Gordillo, P., Knoechel, B., Asmann, Y.W., Slager, S.L., Novak, A.J., Dogan, A., Ansell, S.M., Link, B.K., Zou, L., Gould, J., Saksena, G., Stransky, N., Rangel-Escareño, C., Fernandez-Lopez, J.C., Hidalgo-Miranda, A., Melendez-Zajgla, J., Hernández-Lemus, E., Schwarz-Cruz y Celis, A., Imaz-Rosshandler, I., Ojesina, A.I., Jung, J., Pedamallu, C.S., Lander, E.S., Habermann, T.M., Cerhan, J.R., Shipp, M.A., Getz, G., Golub, T.R.: Discovery and prioritization of somatic mutations in diffuse large B-cell lymphoma (DLBCL) by whole-exome sequencing. Proc. Natl. Acad. Sci. 109, 3879 (2012). 10.1073/pnas.1121343109.

18. Monni, O., Franssila, K., Joensuu, H., Knuutila, S.: BCL2 Overexpression in Diffuse Large B-Cell Lymphoma. Leuk. Lymphoma. 34, 45−52 (1999). 10.3109/10428199909083379.

19. Blenk, S., Engelmann, J., Weniger, M., Schultz, J., Dittrich, M., Rosenwald, A., Müller-Hermelink, H.K., Müller, T., Dandekar, T.: Germinal Center B Cell-Like (GCB) and Activated B Cell-Like (ABC) Type of Diffuse Large B Cell Lymphoma (DLBCL): Analysis of Molecular Predictors, Signatures, Cell Cycle State and Patient Survival. Cancer Inform. 3, 117693510700300004 (2007). 10.1177/117693510700300004.

20. Ahmadi, S.E., Rahimi, S., Zarandi, B., Chegeni, R., Safa, M.: MYC: a multipurpose oncogene with prognostic and therapeutic implications in blood malignancies. J. Hematol. Oncol.J Hematol Oncol. 14, 121 (2021). 10.1186/s13045-021-01111-4.

21. Ott, G., Rosenwald, A., Campo, E.: Understanding MYC-driven aggressive B-cell lymphomas: pathogenesis and classification. Blood. 122, 3884−3891 (2013). 10.1182/blood-2013-05-498329.

22. Slack, G.W., Gascoyne, R.D.: MYC and Aggressive B-cell Lymphomas. Adv. Anat. Pathol. 18, (2011).

23. Sang, Y., Kong, P., Zhang, S., Zhang, L., Cao, Y., Duan, X., Sun, T., Tao, Z., Liu, W.: SGK1 in Human Cancer: Emerging Roles and Mechanisms. Front. Oncol. 10, (2021). 10.3389/fonc.2020.608722.

24. Gao, J., Sidiropoulou, E., Walker, I., Krupka, J.A., Mizielinski, K., Usheva, Z., Samarajiwa, S.A., Hodson, D.J.: SGK1 mutations in DLBCL generate hyperstable protein neoisoforms that promote AKT independence. Blood. 138, 959−964 (2021). 10.1182/blood.2020010432.

25. Merkel, A.L., Meggers, E., Ocker, M.: PIM1 kinase as a target for cancer therapy. Expert Opin. Investig. Drugs. 21, 425−436 (2012). 10.1517/13543784.2012.668527.

26. Brault, L., Menter, T., Obermann, E.C., Knapp, S., Thommen, S., Schwaller, J., Tzankov, A.: PIM kinases are progression markers and emerging therapeutic targets in diffuse large B-cell lymphoma. Br. J. Cancer. 107, 491−500 (2012). 10.1038/bjc.2012.272.

27. Massari, M.E., Rivera, R.R., Voland, J.R., Quong, M.W., Breit, T.M., van Dongen, J.J.M., de Smit, O., Murre, C.: Characterization of ABF-1, a Novel Basic Helix-Loop-Helix Transcription Factor Expressed in Activated B Lymphocytes. Mol. Cell. Biol. 18, 3130 (1998). 10.1128/MCB.18.6.3130.

28. Ushmorov, A., Leithäuser, F., Ritz, O., Barth, T.F.E., Möller, P., Wirth, T.: ABF-1 is frequently silenced by promoter methylation in follicular lymphoma, diffuse large B-cell lymphoma and Burkitt’s lymphoma. Leukemia. 22, 1942 (2008).

29. Galán-Martínez, J., Stamatakis, K., Sánchez-Gómez, I., Vázquez-Cuesta, S., Gironés, N., Fresno, M.: Isoform-specific effects of transcription factor TCFL5 on the pluripotency-related genes SOX2 and KLF4 in colorectal cancer development. Mol. Oncol. 16, 1876−1890 (2022). 10.1002/1878-0261.13085.

30. Errico, M.C., Jin, K., Sukumar, S., Carè, A.: The Widening Sphere of Influence of HOXB7 in Solid Tumors. Cancer Res. 76, 2857−2862 (2016). 10.1158/0008-5472.CAN-15-3444.

31. Nagel, S., Kaufmann, M., Drexler, H.G., MacLeod, R.A.F.: Co-Activation of HOXB7 and BCL2/MYC Via Biallelic IGH Rearrangements in a B-Cell Lymphoma Cell Line. Blood. 104, 4266−4266 (2004). 10.1182/blood.V104.11.4266.4266.

32. Martin, F., Ladoire, S., Mignot, G., Apetoh, L., Ghiringhelli, F.: Human FOXP3 and cancer. Oncogene. 29, 4121−4129 (2010). 10.1038/onc.2010.174.

33. Zhao, Y., Cui, W., Feng, Z., Xue, J., Gulinaer, A., Zhang, W.: Expression of Foxp3 and interleukin-7 receptor and clinicopathological characteristics of patients with diffuse large B-cell lymphoma. Oncol. Lett. 19, 2755−2764 (2020). 10.3892/ol.2020.11374.

34. Chevrier, S., Corcoran, L.M.: BTB-ZF transcription factors, a growing family of regulators of early and late B-cell development. Immunol. Cell Biol. 92, 481−488 (2014). 10.1038/icb.2014.20.

35. Ashburner, M., Ball, C.A., Blake, J.A., Botstein, D., Butler, H., Cherry, J.M., Davis, A.P., Dolinski, K., Dwight, S.S., Eppig, J.T., Harris, M.A., Hill, D.P., Issel-Tarver, L., Kasarskis, A., Lewis, S., Matese, J.C., Richardson, J.E., Ringwald, M., Rubin, G.M., Sherlock, G.: Gene Ontology: tool for the unification of biology. Nat. Genet. 25, 25−29 (2000). 10.1038/75556.

36. Aleksander, S.A., Balhoff, J., Carbon, S., Cherry, J.M., Drabkin, H.J., Ebert, D., Feuermann, M., Gaudet, P., Harris, N.L., Hill, D.P., Lee, R., Mi, H., Moxon, S., Mungall, C.J., Muruganugan, A., Mushayahama, T., Sternberg, P.W., Thomas, P.D., Van Auken, K., Ramsey, J., Siegele, D.A., Chisholm, R.L., Fey, P., Aspromonte, M.C., Nugnes, M.V., Quaglia, F., Tosatto, S., Giglio, M., Nadendla, S., Antonazzo, G., Attrill, H., Dos Santos, G., Marygold, S., Strelets, V., Tabone, C.J., Thurmond, J., Zhou, P., Ahmed, S.H., Asanitthong, P., Luna Buitrago, D., Erdol, M.N., Gage, M.C., Ali Kadhum, M., Li, K.Y.C., Long, M., Michalak, A., Pesala, A., Pritazahra, A., Saverimuttu, S.C.C., Su, R., Thurlow, K.E., Lovering, R.C., Logie, C., Oliferenko, S., Blake, J., Christie, K., Corbani, L., Dolan, M.E., Drabkin, H.J., Hill, D.P., Ni, L., Sitnikov, D., Smith, C., Cuzick, A., Seager, J., Cooper, L., Elser, J., Jaiswal, P., Gupta, P., Jaiswal, P., Naithani, S., Lera-Ramirez, M., Rutherford, K., Wood, V., De Pons, J.L., Dwinell, M.R., Hayman, G.T., Kaldunski, M.L., Kwitek, A.E., Laulederkind, S.J.F., Tutaj, M.A., Vedi, M., Wang, S.-J., D’Eustachio, P., Aimo, L., Axelsen, K., Bridge, A., Hyka-Nouspikel, N., Morgat, A., Aleksander, S.A., Cherry, J.M., Engel, S.R., Karra, K., Miyasato, S.R., Nash, R.S., Skrzypek, M.S., Weng, S., Wong, E.D., Bakker, E., Berardini, T.Z., Reiser, L., Auchincloss, A., Axelsen, K., Argoud-Puy, G., Blatter, M.C., Boutet, E., Breuza, L., Bridge, A., Casals-Casas, C., Coudert, E., Estreicher, A., Livia Famiglietti, M., Feuermann, M., Gos, A., Gruaz-Gumowski, N., Hulo, C., Hyka-Nouspikel, N., Jungo, F., Le Mercier, P., Lieberherr, D., Masson, P., Morgat, A., Pedruzzi, I., Pourcel, L., Poux, S., Rivoire, C., Sundaram, S., Bateman, A., Bowler-Barnett, E., Bye-A-Jee, H., Denny, P., Ignatchenko, A., Ishtiaq, R., Lock, A., Lussi, Y., Magrane, M., Martin, M.J., Orchard, S., Raposo, P., Speretta, E., Tyagi, N., Warner, K., Zaru, R., Diehl, A.D., Lee, R., Chan, J., Diamantakis, S., Raciti, D., Zarowiecki, M., Fisher, M., James-Zorn, C., Ponferrada, V., Zorn, A., Ramachandran, S., Ruzicka, L., Westerfield, M.: The Gene Ontology knowledgebase in 2023. Genetics. 224, (2023). 10.1093/genetics/iyad031.

37. Luck, K., Kim, D.-K., Lambourne, L., Spirohn, K., Begg, B.E., Bian, W., Brignall, R., Cafarelli, T., Campos-Laborie, F.J., Charloteaux, B., Choi, D., Coté, A.G., Daley, M., Deimling, S., Desbuleux, A., Dricot, A., Gebbia, M., Hardy, M.F., Kishore, N., Knapp, J.J., Kovács, I.A., Lemmens, I., Mee, M.W., Mellor, J.C., Pollis, C., Pons, C., Richardson, A.D., Schlabach, S., Teeking, B., Yadav, A., Babor, M., Balcha, D., Basha, O., Bowman-Colin, C., Chin, S.-F., Choi, S.G., Colabella, C., Coppin, G., D’Amata, C., De Ridder, D., De Rouck, S., Duran-Frigola, M., Ennajdaoui, H., Goebels, F., Goehring, L., Gopal, A., Haddad, G., Hatchi, E., Helmy, M., Jacob, Y., Kassa, Y., Landini, S., Li, R., van Lieshout, N., MacWilliams, A., Markey, D., Paulson, J.N., Rangarajan, S., Rasla, J., Rayhan, A., Rolland, T., San-Miguel, A., Shen, Y., Sheykhkarimli, D., Sheynkman, G.M., Simonovsky, E., Taşan, M., Tejeda, A., Tropepe, V., Twizere, J.-C., Wang, Y., Weatheritt, R.J., Weile, J., Xia, Y., Yang, X., Yeger-Lotem, E., Zhong, Q., Aloy, P., Bader, G.D., De Las Rivas, J., Gaudet, S., Hao, T., Rak, J., Tavernier, J., Hill, D.E., Vidal, M., Roth, F.P., Calderwood, M.A.: A reference map of the human binary protein interactome. Nature. 580, 402−408 (2020). 10.1038/s41586-020-2188-x.

38. Grover, A., Leskovec, J.: Node2vec: Scalable Feature Learning for Networks. In: Proceedings of the 22nd ACM SIGKDD International Conference on Knowledge Discovery and Data Mining. pp. 855−864. Association for Computing Machinery, New York, NY, USA (2016). 10.1145/2939672.2939754.

39. Ke, G., Meng, Q., Finley, T., Wang, T., Chen, W., Ma, W., Ye, Q., Liu, T.-Y.: LightGBM: A Highly Efficient Gradient Boosting Decision Tree. In: Guyon, I., Luxburg, U.V., Bengio, S., Wallach, H., Fergus, R., Vishwanathan, S., and Garnett, R. (eds.) Advances in Neural Information Processing Systems. Curran Associates, Inc. (2017).

40. Zhang, J., Bajari, R., Andric, D., Gerthoffert, F., Lepsa, A., Nahal-Bose, H., Stein, L.D., Ferretti, V.: The International Cancer Genome Consortium Data Portal. Nat. Biotechnol. 37, 367−369 (2019). 10.1038/s41587-019-0055-9.

41. Kent, W.J., Sugnet, C.W., Furey, T.S., Roskin, K.M., Pringle, T.H., Zahler, A.M., Haussler, and D.: The Human Genome Browser at UCSC. Genome Res. 12, 996−1006 (2002). 10.1101/gr.229102.

42. Tate, J.G., Bamford, S., Jubb, H.C., Sondka, Z., Beare, D.M., Bindal, N., Boutselakis, H., Cole, C.G., Creatore, C., Dawson, E., Fish, P., Harsha, B., Hathaway, C., Jupe, S.C., Kok, C.Y., Noble, K., Ponting, L., Ramshaw, C.C., Rye, C.E., Speedy, H.E., Stefancsik, R., Thompson, S.L., Wang, S., Ward, S., Campbell, P.J., Forbes, S.A.: COSMIC: the Catalogue Of Somatic Mutations In Cancer. Nucleic Acids Res. gky1015−gky1015 (2018). 10.1093/nar/gky1015.

43. Mathelier, A., Zhao, X., Zhang, A.W., Parcy, F., Worsley-Hunt, R., Arenillas, D.J., Bucman, S., Chen, C., Chou, A., Ienasescu, H., Lim, J., Shyr, C., Tan, G., Zhou, M., Lenhard, B., Sandelin, A., Wasserman, W.W.: JASPAR 2014: an extensively expanded and updated open-access database of transcription factor binding profiles. Nucleic Acids Res. 42, D142− D147 (2013). 10.1093/nar/gkt997.

44. Lonsdale, J., Thomas, J., Salvatore, M., Phillips, R., Lo, E., Shad, S., Hasz, R., Walters, G., Garcia, F., Young, N., Foster, B., Moser, M., Karasik, E., Gillard, B., Ramsey, K., Sullivan, S., Bridge, J., Magazine, H., Syron, J., Fleming, J., Siminoff, L., Traino, H., Mosavel, M., Barker, L., Jewell, S., Rohrer, D., Maxim, D., Filkins, D., Harbach, P., Cortadillo, E., Berghuis, B., Turner, L., Hudson, E., Feenstra, K., Sobin, L., Robb, J., Branton, P., Korzeniewski, G., Shive, C., Tabor, D., Qi, L., Groch, K., Nampally, S., Buia, S., Zimmerman, A., Smith, A., Burges, R., Robinson, K., Valentino, K., Bradbury, D., Cosentino, M., Diaz-Mayoral, N., Kennedy, M., Engel, T., Williams, P., Erickson, K., Ardlie, K., Winckler, W., Getz, G., DeLuca, D., MacArthur, D., Kellis, M., Thomson, A., Young, T., Gelfand, E., Donovan, M., Meng, Y., Grant, G., Mash, D., Marcus, Y., Basile, M., Liu, J., Zhu, J., Tu, Z., Cox, N.J., Nicolae, D.L., Gamazon, E.R., Im, H.K., Konkashbaev, A., Pritchard, J., Stevens, M., Flutre, T., Wen, X., Dermitzakis, E.T., Lappalainen, T., Guigo, R., Monlong, J., Sammeth, M., Koller, D., Battle, A., Mostafavi, S., McCarthy, M., Rivas, M., Maller, J., Rusyn, I., Nobel, A., Wright, F., Shabalin, A., Feolo, M., Sharopova, N., Sturcke, A., Paschal, J., Anderson, J.M., Wilder, E.L., Derr, L.K., Green, E.D., Struewing, J.P., Temple, G., Volpi, S., Boyer, J.T., Thomson, E.J., Guyer, M.S., Ng, C., Abdallah, A., Colantuoni, D., Insel, T.R., Koester, S.E., Little, A.R., Bender, P.K., Lehner, T., Yao, Y., Compton, C.C., Vaught, J.B., Sawyer, S., Lockhart, N.C., Demchok, J., Moore, H.F.: The Genotype-Tissue Expression (GTEx) project. Nat. Genet. 45, 580−585 (2013). 10.1038/ng.2653.

45. Ashburner, M., Ball, C.A., Blake, J.A., Botstein, D., Butler, H., Cherry, J.M., Davis, A.P., Dolinski, K., Dwight, S.S., Eppig, J.T., Harris, M.A., Hill, D.P., Issel-Tarver, L., Kasarskis, A., Lewis, S., Matese, J.C., Richardson, J.E., Ringwald, M., Rubin, G.M., Sherlock, G.: Gene ontology: tool for the unification of biology. The Gene Ontology Consortium. Nat. Genet. 25, 25−29 (2000). 10.1038/75556.

