## Supplementary information for "Novel Driver Mutations in GCB Lymphoma Patients That Affect Transcription Factors Binding"

### Supplementary Data

#### 1 Machine learning results data

**Table 1.** Grid search results of different parameters (Node2Vec vector length, Node2Vec walk length, and Node2Vec window size) and different models (GO alone, PPI alone, merged GO and PPI and a reference GO model with randomized nodes). The results are represented by accuracy and were calculated using 10-fold cross-validation. The best-performing model for each parameter set is highlighted in bold.

| Node2Vec length | Walk length | Window size | GO accuracy | PPI accuracy | Merged PPI+GO accuracy | Randomized GO accuracy |
|---|---|---|---|---|---|---|
| 8 | 4 | 4 | 0.559 | 0.499 | **0.569** | 0.498 |
| 8 | 4 | 8 | 0.538 | 0.503 | **0.549** | 0.484 |
| **8** | **4** | **16** | **0.589** | 0.515 | 0.569 | 0.492 |
| 8 | 8 | 4 | 0.548 | 0.531 | **0.552** | 0.487 |
| 8 | 8 | 8 | **0.551** | 0.464 | 0.544 | 0.527 |
| 8 | 8 | 16 | **0.543** | 0.484 | **0.543** | 0.49 |
| 8 | 16 | 4 | **0.584** | 0.478 | 0.547 | 0.462 |
| 8 | 16 | 8 | 0.566 | 0.507 | **0.582** | 0.463 |
| 8 | 16 | 16 | **0.55** | 0.482 | 0.536 | 0.502 |
| 16 | 4 | 4 | **0.553** | 0.475 | 0.539 | 0.452 |
| 16 | 4 | 8 | **0.543** | 0.477 | 0.514 | 0.469 |
| 16 | 4 | 16 | **0.572** | 0.51 | 0.556 | 0.485 |
| 16 | 8 | 4 | **0.541** | 0.528 | 0.518 | 0.479 |
| 16 | 8 | 8 | 0.539 | 0.466 | 0.522 | **0.541** |
| 16 | 8 | 16 | 0.53 | 0.534 | **0.54** | 0.499 |
| 16 | 16 | 4 | **0.581** | 0.52 | **0.581** | 0.474 |
| 16 | 16 | 8 | **0.557** | 0.492 | 0.527 | 0.475 |
| 16 | 16 | 16 | **0.559** | 0.476 | 0.522 | 0.476 |
| 32 | 4 | 4 | **0.526** | 0.483 | 0.511 | 0.5 |
| 32 | 4 | 8 | **0.556** | 0.478 | 0.512 | 0.532 |
| 32 | 4 | 16 | **0.529** | 0.502 | 0.513 | 0.508 |
| 32 | 8 | 4 | **0.541** | 0.512 | 0.535 | 0.473 |
| 32 | 8 | 8 | **0.552** | 0.489 | 0.517 | 0.479 |
| 32 | 8 | 16 | **0.534** | 0.494 | 0.513 | 0.493 |
| 32 | 16 | 4 | **0.534** | 0.489 | 0.498 | 0.495 |
| 32 | 16 | 8 | 0.531 | 0.5 | 0.516 | **0.56** |
| 32 | 16 | 16 | **0.569** | 0.495 | 0.553 | 0.507 |

#### 2 Additional mutations blocks data

Following this, detailed results of five additional mutations blocks that may affect MSC binding will be presented as examples of the pipeline's results. Among the 56 identified mutation blocks, six may be associated with MSC binding. All six mutations appear to cause the upregulation of proto-oncogenes involved in apoptosis and cell development.

**Mutations block #1 – SGK1**

This mutations block includes six different missense mutations that were found 12 times (out of 241 DLBCL patients). The block is located on chromosome 6: 134174583- 134174596, where the SGK1 gene is found, see Fig. 1. SGK1 is a known oncogene and its overexpression is correlated with a wide range of cancers, including DLBCL[1, 2].

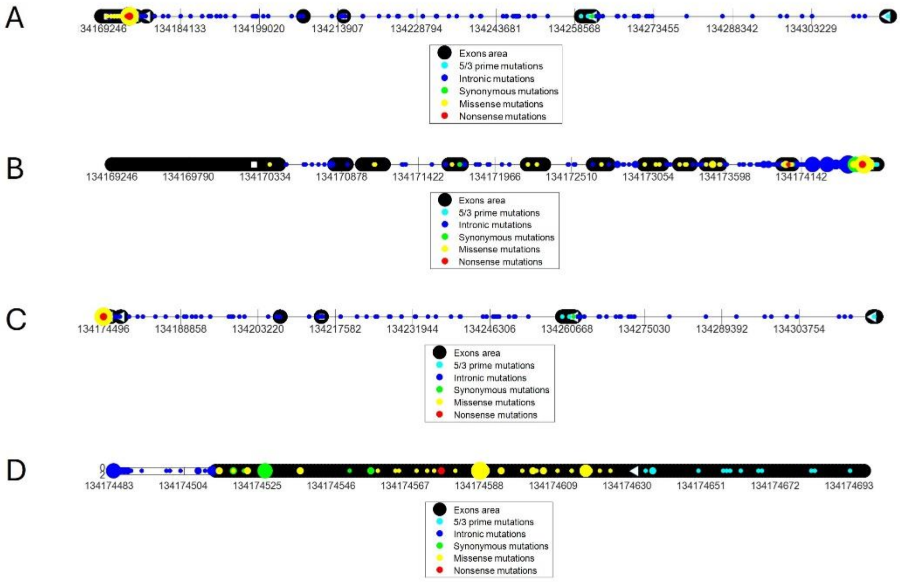

**Fig 1.** This figure illustrates the distribution of mutations in SGK1 among malignant lymphoma patients along the SGK1 gene while the size of each circle represents the frequency of a given mutation in patients. The figure is divided into four parts, where A shows the mutations along the entire gene. B shows the mutations from the beginning of the gene up until the mutation block in 134174583- 134174596. C shows the mutations from the mutations block in 134174583- 134174596 up until the end of the gene. D shows the mutations in the area of the mutation block (+- 100 nucleotides). As can be seen the 134174583- 134174596 mutations block contains several missense mutations, one of which is highly recurrent.

To validate the effect of this mutation on the gene, its expression was analyzed. The expression of the SGK1 gene in patients with the mutation was higher than in randomized groups of DLBCL patients (Pvalue = 0.0089, see Fig. 2), Which aligns with previous findings that SGK1 overexpression is correlated with DLBCL[2].

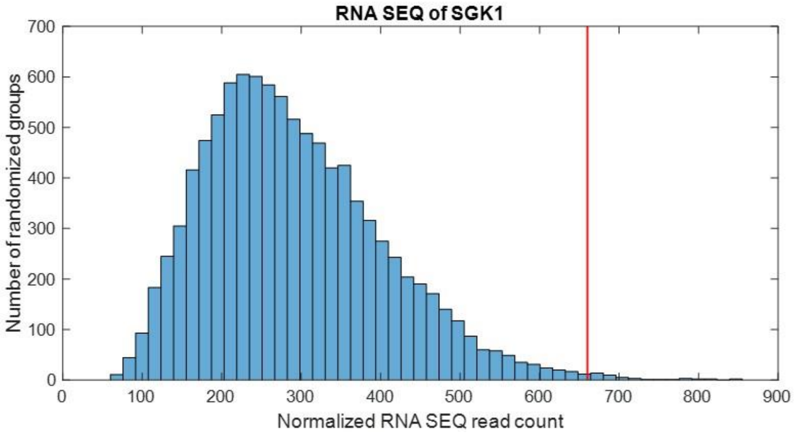


**Fig 2.** Using the RNA sequence data, 10,000 groups of patients were randomly selected, and their mean expression levels were calculated. The red line represents the mean expression of patients with mutations in the 134174583-134174596 block of the SGK1 gene.

A search was conducted for relevant transcription factors that might explain the change in expression. Only transcription factors expressed in EBV-transformed lymphocytes were considered and were examined for changes in their binding ability to the relevant region. Only two transcription factors were identified as potential contributors to the observed overexpression:

*Transcription factor candidate #1 – MSC:*

The mutations in the mutations blocks interfere with the ability of the MSC repressor to bind to the forward strand at this location, as shown in Fig. 3. The reduced binding ability of the repressor may explain the gene's upregulation.

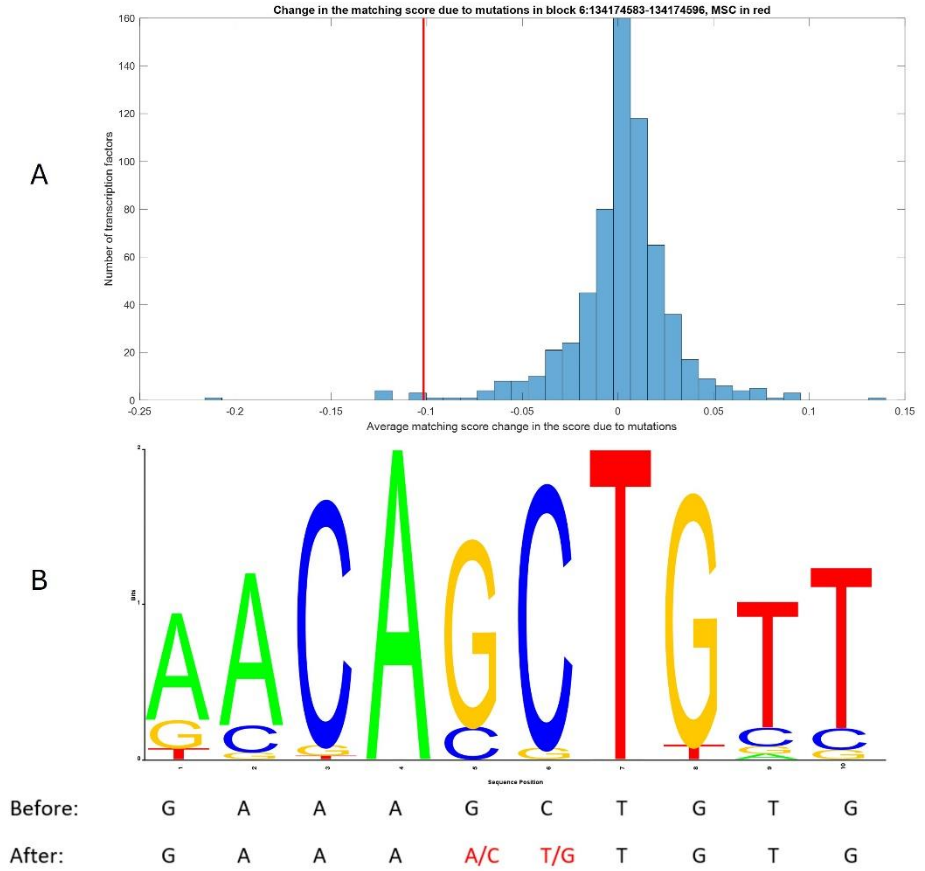


**Fig 3.** Represents the effect of mutations on the ability of transcription factors to bind to the gene. In A, a histogram representing the influence of the mutation in the block on the ability of each transcription factor to bind. As can be seen, most transcription factors are only minimally affected by these mutations, while a few show significant changes. MSC is indicated by a red line. In B, The LOGO of the binding site of MSC according to JASPAR’s PSSMs. Below, the relevant nucleotide sequence (6: 134174582- 134174591 forward strand) is shown with the mutation (after) and without the mutation (before). Marked in red are alternatives nucleotides that were found in patients.

*Transcription factor candidate #2 – ASCL1:*

The mutations in the mutations blocks interfere with the ability of the ASCL1 repressor to bind to the reverse strand of this location, as shown in Fig 4. ASCL1 has been implicated in various cancers, including lung cancer [3] and leukemia[4]. The reduction in the repressor's binding ability to the genome may help explain the upregulation of the gene.

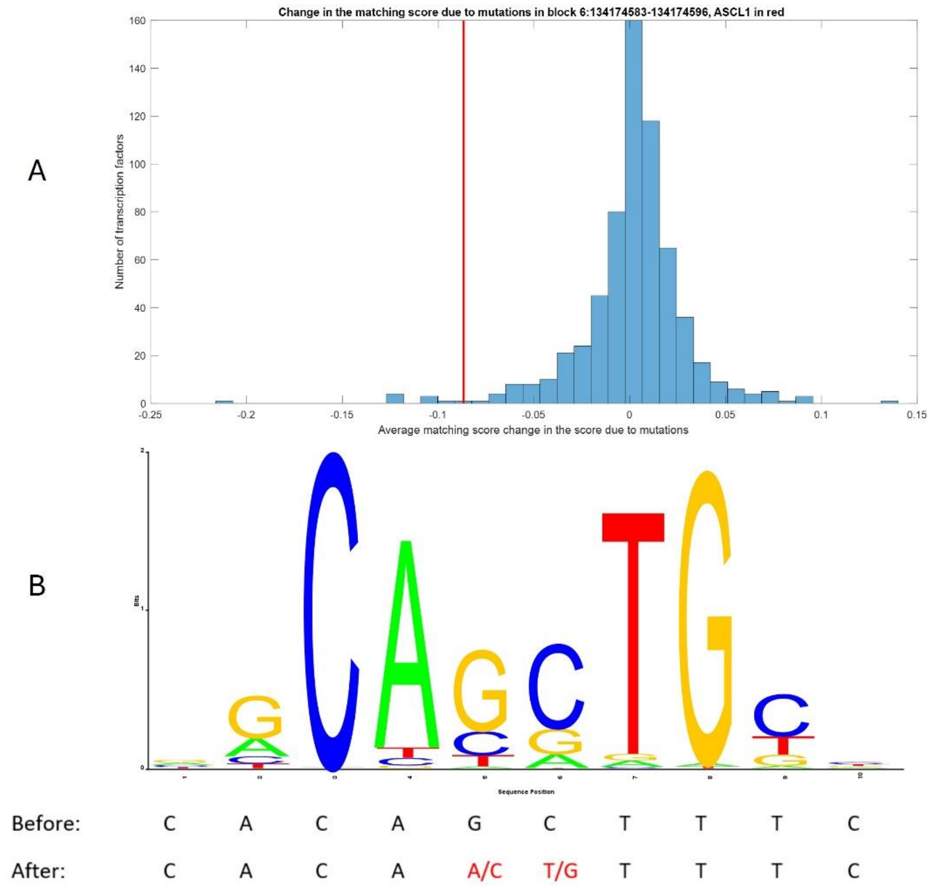


**Fig 4.** Represents the effect the mutations have on the ability of transcription factors to bind to the gene. In A, a histogram representing the influence of the mutation in the block on the ability of each transcription factor to bind. As can be seen, most transcription factors are minimally affected by this mutation, while only a few are highly effected. ASCL1 is marked in red line. In B, the LOGO of the binding site of ASCL1 according to JASPAR's PSSMs. Below the nucleotides of the relevant site (6: 134174582- 134174591 reverse strand) is shown with the mutation (after) and without the mutation (before). Marked in red are alternatives nucleotides that were found in patients.

**Mutations block #2 – MYC (127736580)**

This mutations block includes eleven different 5' prime mutations that was found 14 times (out of 241 DLBCL patients). The block is located on chromosome 8: 127736580- 127736592, Where the gene MYC can be found (see Fig 5). "MYC is a known oncogene, and its overexpression has been linked to DLBCL[5].

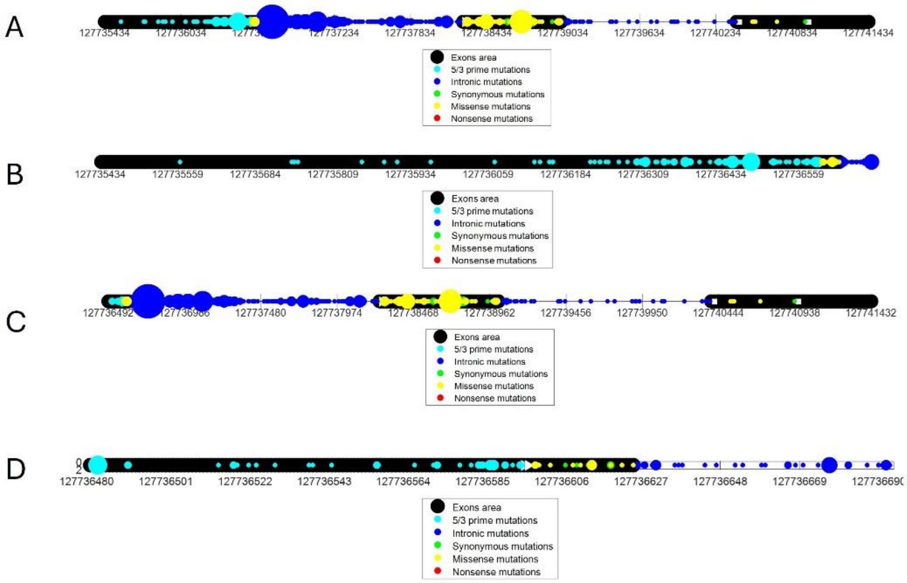


**Fig 5.** This figure illustrates the distribution of mutations in the MYC gene among DLBCL patients along the MYC gene while the size of each circle represents the frequency of a given mutation among patients. The figure is divided into four parts, where A shows the mutations along the entire gene. B shows the mutations from the beginning of the gene up until the mutation block in 127736580- 127736592. C shows the mutations from the mutations block in 127736580-127736592 up until the end of the gene. D shows the mutations in the area of the mutation block (+- 100 nucleotides). As can be seen the 127736580- 127736592 mutations block includes several distinct 5′-prime mutations.

To validate the effect of this mutation on gene expression, the expression levels of the MYC gene were analyzed. The expression of the MYC gene in patients with the mutation was higher than in randomized groups of DLBCL patients (Pvalue = 0.0227, see Fig. 6). This overexpression of MYC may contribute to cancer progression, given that MYC is a well-established driver of cellular growth.

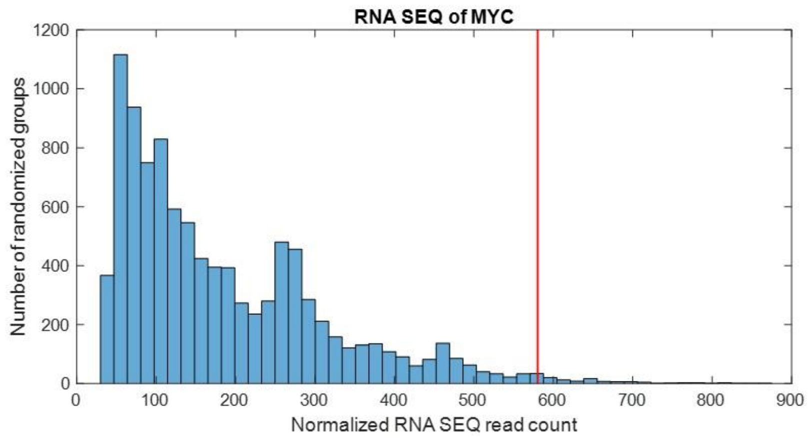


**Fig 6.** Using the RNA sequence data, 10,000 groups of patients were randomly selected, and their mean expression was calculated. The red line represents the mean expression of patients with mutations in the 127736580- 127736592 block in the MYC gene.

Now a search for relevant transcription factors that might explain the change in the expression was conducted. Only transcription factors that are expressed in EBV transformed lymphocytes were considered and were assessed for changes in their binding ability to the relevant area. Only two transcription factors were found as a possible cause to the aforementioned overexpression:

*Transcription factor candidate #1 – MSC:*

The mutations in the mutations blocks interfere with the ability of the MSC repressor to bind to the forward strand of this location, as shown in Fig. 7. The reduction in the repressor's ability to bind to the genome may explain the upregulation of the gene.

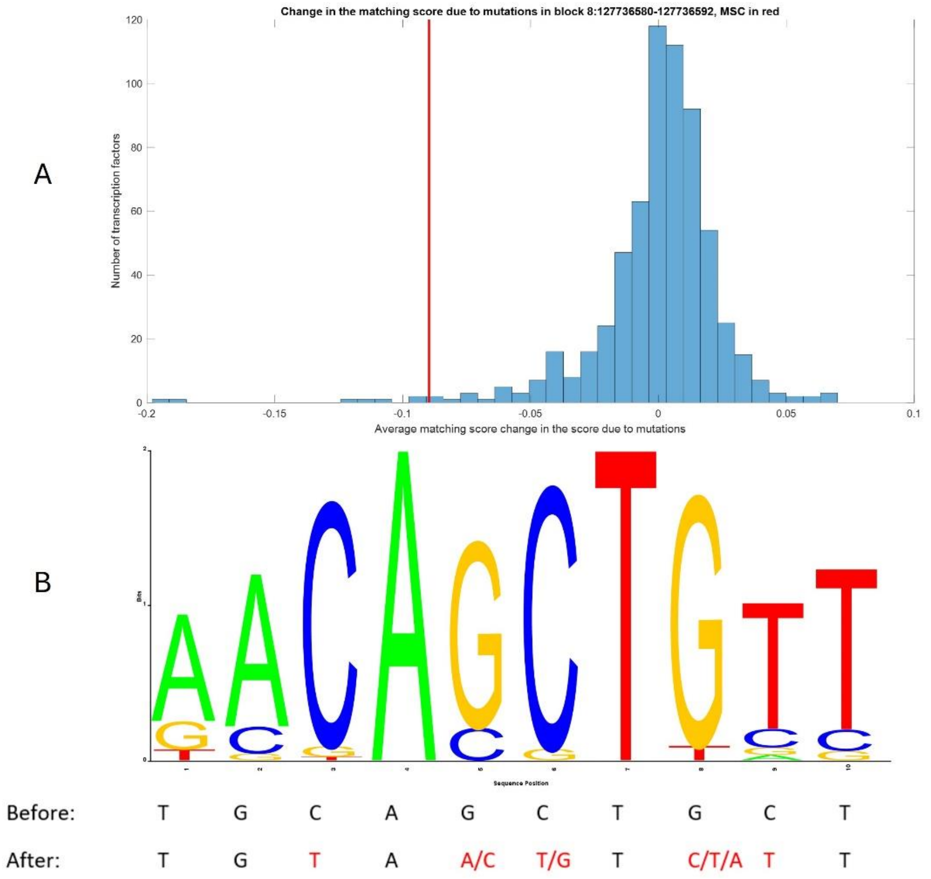


**Fig 7.** Represents the effect of mutations on the ability of transcription factors to bind to the gene. In A, a histogram representing the influence of the mutation in the block on the ability of each transcription factor to bind. As can be seen most transcription factors are minimally affected by this mutation, while only a few are highly affected. MSC is marked in red line. In B, The LOGO of the binding site of MSC according to JASPAR's PSSMs. Below the nucleotides of the relevant site (6: 127736578- 127736587 forward strand) is shown with the mutation (after) and without the mutation (before). Marked in red are alternatives nucleotides that were found in patients.

*Transcription factor candidate #2 – ASCL1:*

The mutations in the mutations blocks interfere with the ability of the ASCL1 repressor to bind to the reverse strand of this location, as shown in Fig. 8. The decrease in the ability of a repressor to bind to the genome may explain the upregulation of the gene.

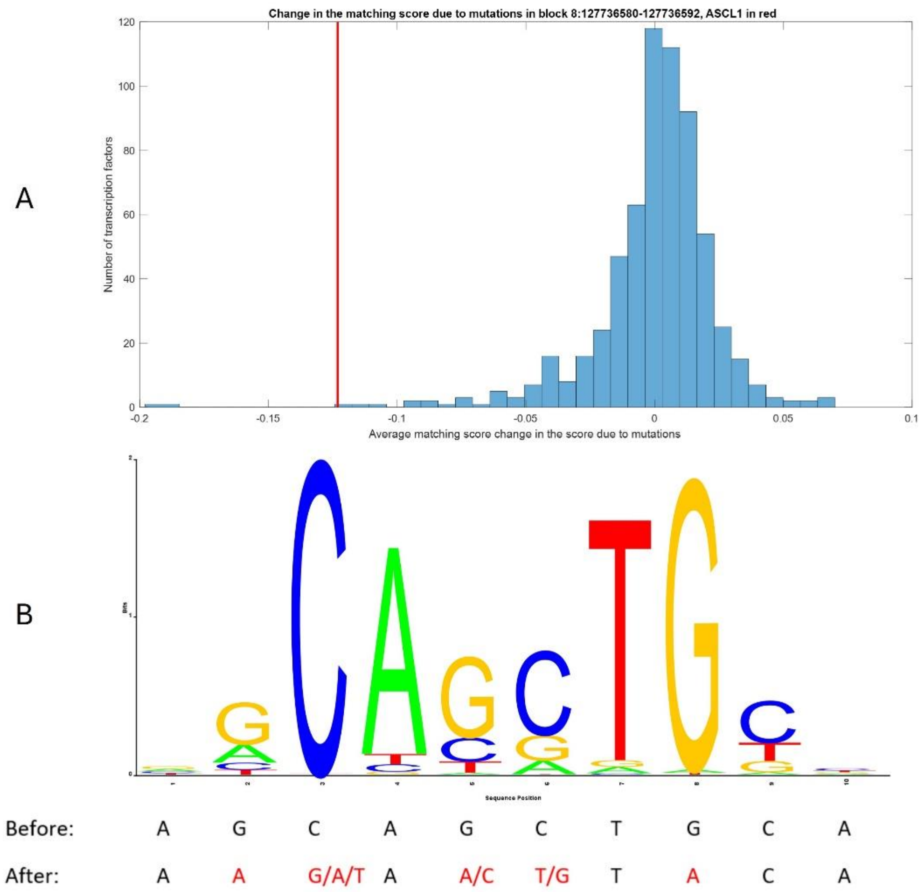


**Fig 8.** Represents the effect the mutations have on the ability of transcription factors to bind to the gene. In A, a histogram representing the influence of the mutation in the block on the ability of each transcription factor to bind. As can be seen, most transcription factors are minimally affected by this mutation, while only a few are highly effected. ASCL1 is marked in red line. In B, The LOGO of the binding site of ASCL1 according to JASPAR's PSSMs. Below the nucleotides of the relevant site (6: 127736578- 127736587 reverse strand) is shown with the mutation (after) and without the mutation (before). Marked in red are alternatives nucleotides that were found in patients.

#### Mutations block #3 – MYC (127738131)

This mutations block includes eight different intron mutations that were found 9 times (out of 241 DLBCL patients). The block is located on chromosome 8: 127738131-127738141, where the gene MYC can be found (see Fig. 9).

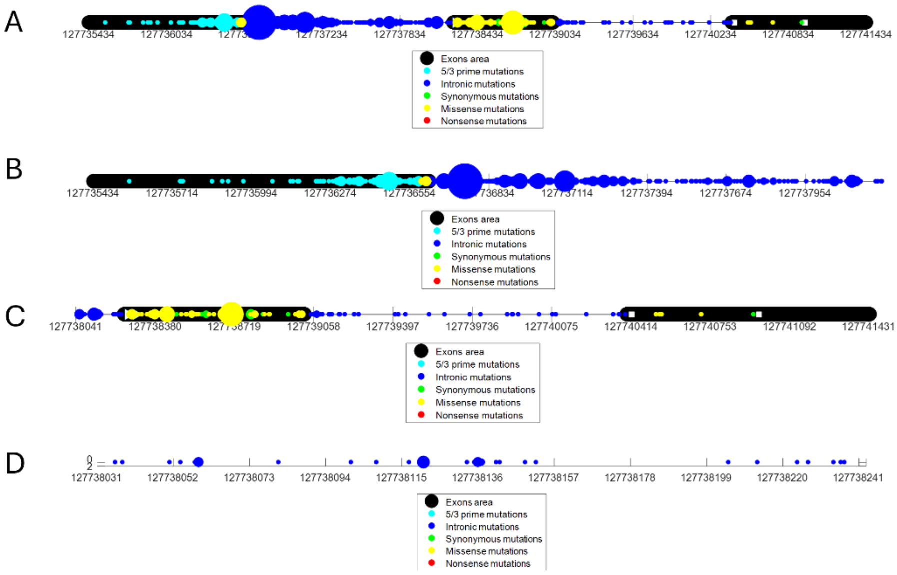

**Fig 9.** This figure represents the locations of mutations in DLBCL patients along the MYC gene, with the size of each circle indicating the frequency of mutations in patients. The figure is divided into four parts, where A shows the mutations along the entire gene. B shows the mutations from the beginning of the gene up until the mutation block in 127738131- 127738141. C shows the mutations from the mutations block in 127738131- 127738141 up until the end of the gene. D shows the mutations in the area of the mutation block (+- 100 nucleotides). As can be seen the 127738131- 127738141 mutations block contains several different intron mutations.

To validate that this mutation has an effect on the gene, its expression was assessed. The expression of the MYC gene in patients with the mutation was higher compared to randomized groups of malignant lymphoma patients (Pvalue = 0.0247, see Fig. 10). This overexpression of MYC may provide the cancer with a growth advantage, as this is growth promoting gene.

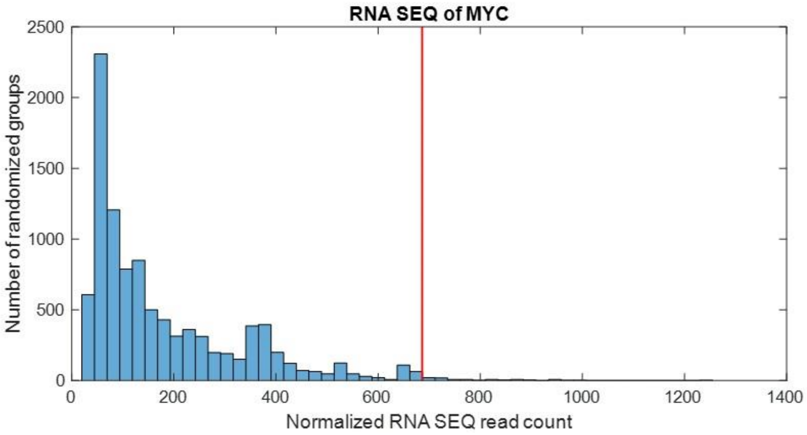


**Fig 10.** Using the RNA sequencing data, 10,000 random patient groups were selected, and their mean expression was calculated. The red line represents the mean expression of patients with mutations in the 127738131- 127738141 block in MYC gene.

Now a search for relevant transcription factors that might explain the change in the expression was conducted. Only transcription factors that are expressed in EBV transformed lymphocytes were considered and were checked for a change in the binding ability of the transcription factor to the relevant area. Only three transcription factors were found as a possible cause to the mentioned-above overexpression:

*Transcription factor candidate #1 – MSC:*

The mutations in the mutations blocks interfere with the ability of the MSC repressor to bind to the forward strand of this location, as shown in Fig. 11. The decrease in the repressor's ability to bind to the genome might explain the upregulation of the gene.

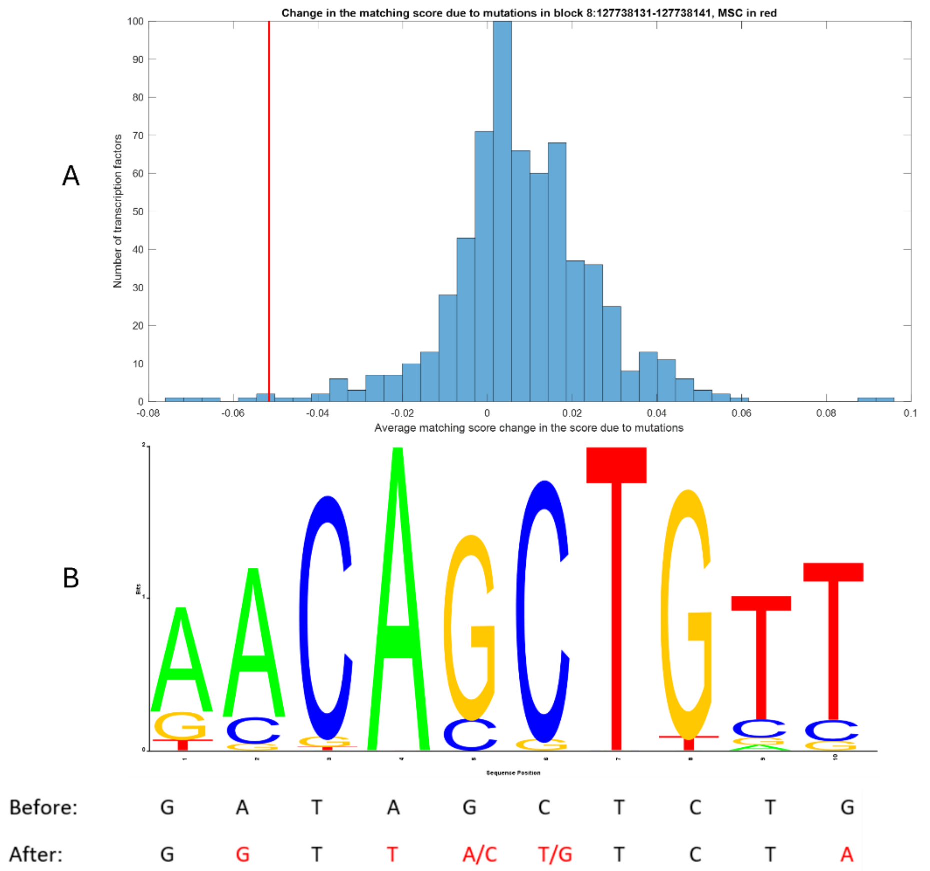


**Fig 11.** Represents the effect of the mutations on the ability of transcription factors to bind to the gene. In A, a histogram representing the influence of the mutation in the block on the ability of each transcription factor to bind. As can be seen, most transcription factors are minimally affected by this mutation, while only a few are highly effected. MSC is marked in red line. In B The LOGO of the binding site of MSC according to JASPAR's PSSMs. Below the nucleotides of the relevant site (8: 127738130- 127738139 forward strand) is shown with the mutation (after) and without the mutation (before). Marked in red are alternatives nucleotides that were found in patients.

*Transcription factor candidate #2 – SOX9:*

The mutations in the mutations blocks increase the ability of the SOX9 activator to bind to the forward strand of this location, as shown in Fig. 12. SOX9 plays a role in various types of cancer[6] and it was found to be overexpressed in DLBCL[7]. The increased ability of the activator to bind to the genome might explain the upregulation of the MYC gene.

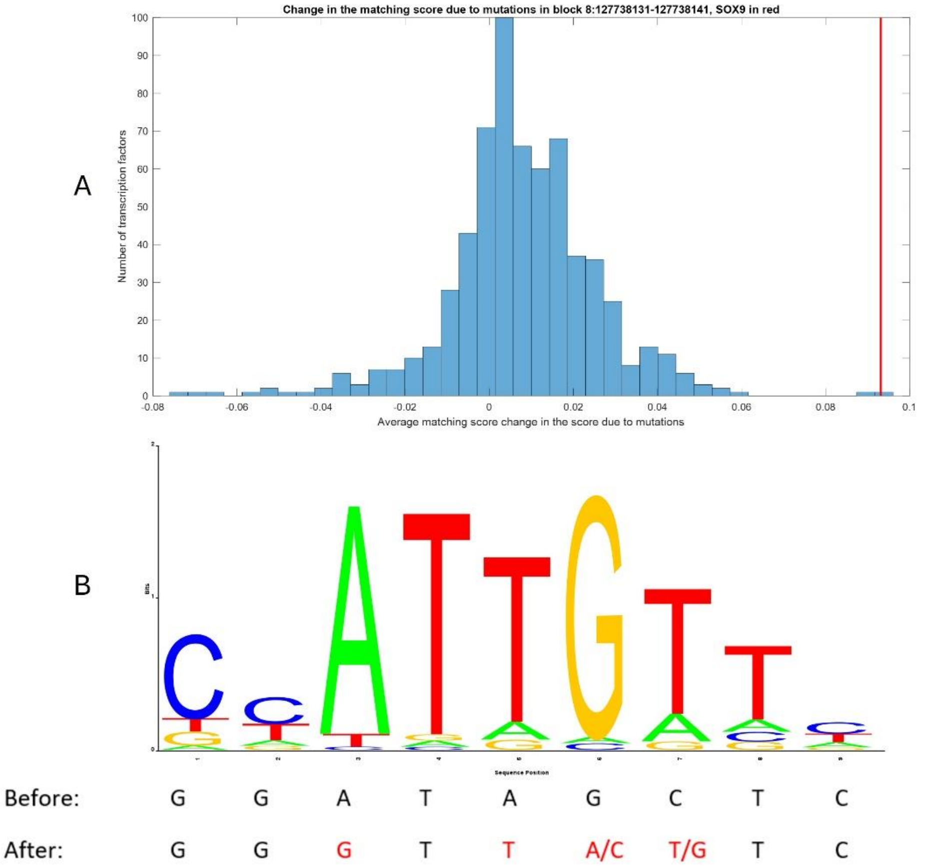


**Fig 12.** Represents the effect of the mutations on the ability of transcription factors to bind to the gene. In A, a histogram representing the influence of the mutation in the block on the ability of each transcription factor to bind. As can be seen, most transcription factors are minimally affected by this mutation, while only a few are highly affected. SOX9 is marked with a red line. In B, The LOGO of the binding site of SOX9 according to JASPAR's PSSMs. Below the nucleotides of the relevant site (8: 127738129-127738137 forward strand) is shown with the mutation (after) and without the mutation (before). Marked in red are alternatives nucleotides that were found in patients.

*Transcription factor candidate #3 – YY1*

The mutations in the mutations blocks interfere with the ability of the YY1 repressor to bind to the forward strand of this location, as shown in Fig. 13. YY1 regulates the cell cycle and plays a significant role in cancer diseases[8] and DLBCL specifically[9]. The decreased ability of this repressor to bind to the genome may explain the upregulation of the MYC gene.

A

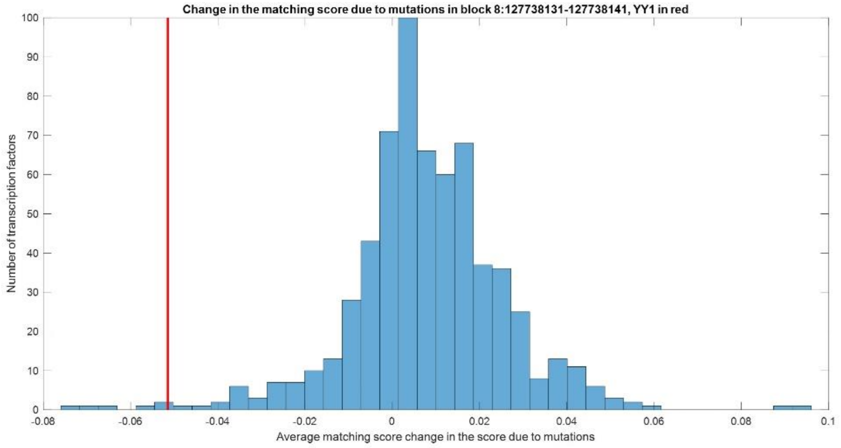


B

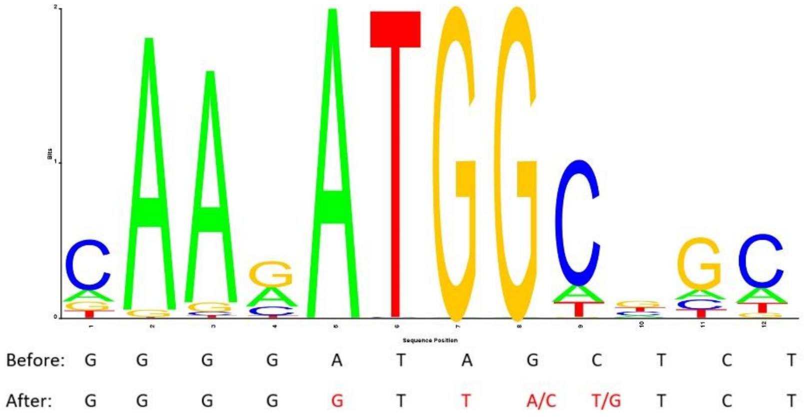


**Fig 13.** Represents the effect the mutations have on the ability of transcription factors to bind to the gene. In A, a histogram representing the influence of the mutation in the block on the ability of each transcription factor to bind. As can be seen, most transcription factors are minimally affected by this mutation, while only a few are highly affected. YY1 is marked in red line. In B, The LOGO of the binding site of SOX9 according to JASPAR's PSSMs. Below the nucleotides of the relevant site (8: 127738127- 127738138 forward strand) is shown with the mutation (after) and without the mutation (before). Marked in red are alternatives nucleotides that were found in patients.

**Mutations block #4 – BCL2 (63318598)**

This mutations block includes eight different missense and synonymous mutations that were found 32 times (out of 241 DLBCL patients). The block is located on chromosome 18: 63318598- 63318610, where the gene BCL2 is found and some of these mutations were previously suspected to be related to transcription factors[10] (see Fig. 14). BCL2 is a known oncogene that prevent apoptosis. Upregulation of BCL2 was corelated with DLBCL[11, 12]

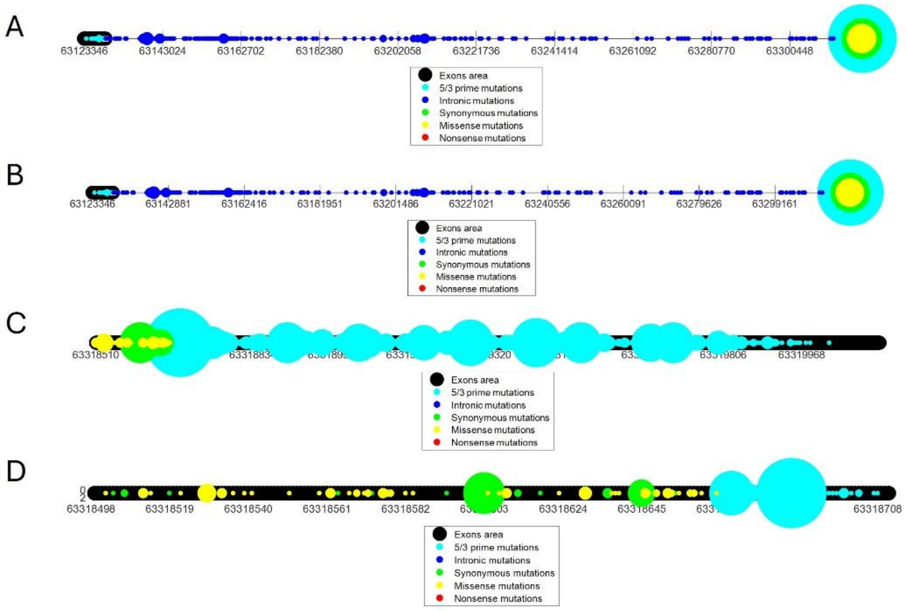


**Fig 14.** This figure represents where mutations of DLBCL patients can be found along the BCL2 gene while the size of each circle represents the frequency of the mutation in patients. The figure is divided into four parts, where A shows the mutations along the entire gene. B shows the mutations from the beginning of the gene up until the mutation block in 63318598- 63318610. C shows the mutations from the mutations block in 63318598- 63318610 up until the end of the gene. D shows the mutations in the area of the mutation block (+- 100 nucleotides). As can be seen the 63318598- 63318610 mutations block includes different missense and synonymous mutations where some synonymous mutations are highly recurrent.

To validate that this mutation has an effect on the gene, the expression was checked. The expression of the BCL2 gene in patients with the mutation was higher than in randomized groups of malignant lymphoma patients (Pvalue = 0.0199, see Fig. 15). This overexpression of BCL2 may contribute to cancer progression as this gene prevents apoptosis.

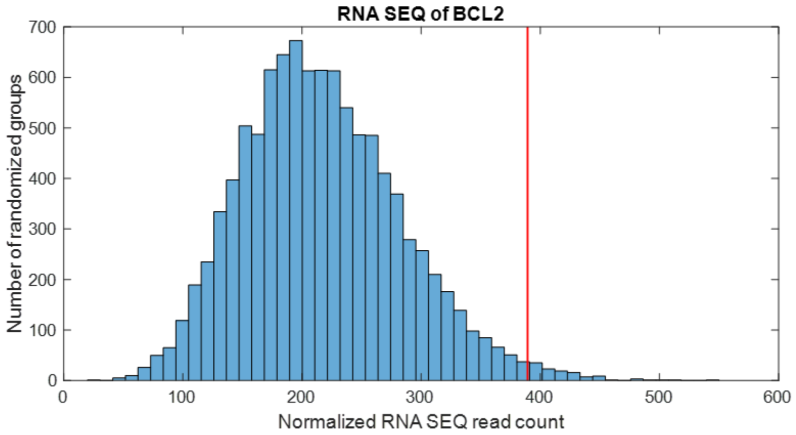


**Fig 15.** Using the RNA sequence data, 10,000 groups of patients were randomly selected, and their mean expression was calculated. The red line represents the mean expression of patients with mutations in the 63318598- 63318610 block in the BCL2 gene.

Now a search for relevant transcription factors that might explain the change in the expression was conducted. Only transcription factors that are expressed in EBV transformed lymphocytes were considered and were checked for a change in the binding ability of the transcription factor to the relevant area. Only one transcription factor was found as a possible cause to the mentioned-above overexpression – MSC. The mutations in the mutations block interfere with the binding of the MSC repressor to BCL2, as shown in Fig. 16, which may lead to the overexpression of BCL2.

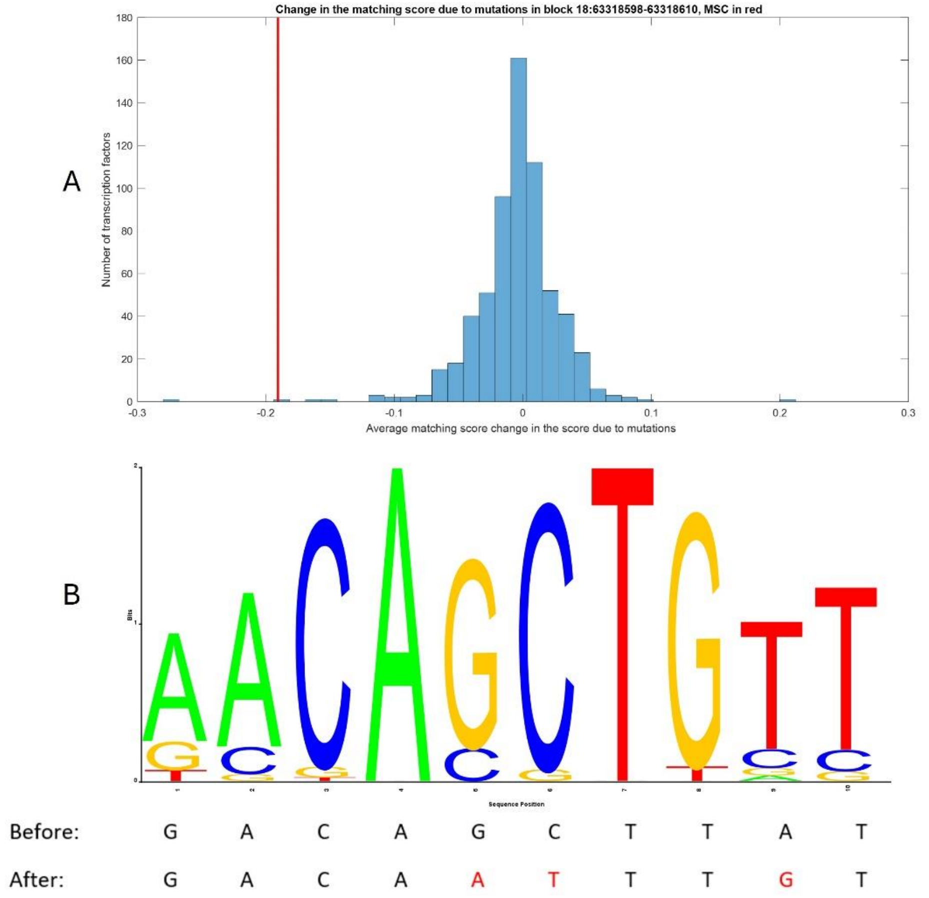


**Fig 16.** Represents the effect of the mutations on the ability of transcription factors to bind to the gene. In A, a histogram representing the influence of the mutation in the block on the ability of each transcription factor to bind. As can be seen, most transcription factors are minimally affected by this mutation, while only a few are highly effected. MSC is marked with a red line. In B, The LOGO of the binding site of MSC according to JASPAR's PSSMs. Below the nucleotides of the relevant site (18: 63318596- 63318605 forward strand) is shown with the mutation (after) and without the mutation (before). Marked in red are alternatives nucleotides that were found in patients.

*Mutations block #5 – BCL2 (63319276)*

This mutations block includes 14 different 5' prime mutations, found 39 times (out of 241 DLBCL patients). The block is located on chromosome 18: 63319276-63319286, where the gene BCL2 can been found (see Fig. 17).

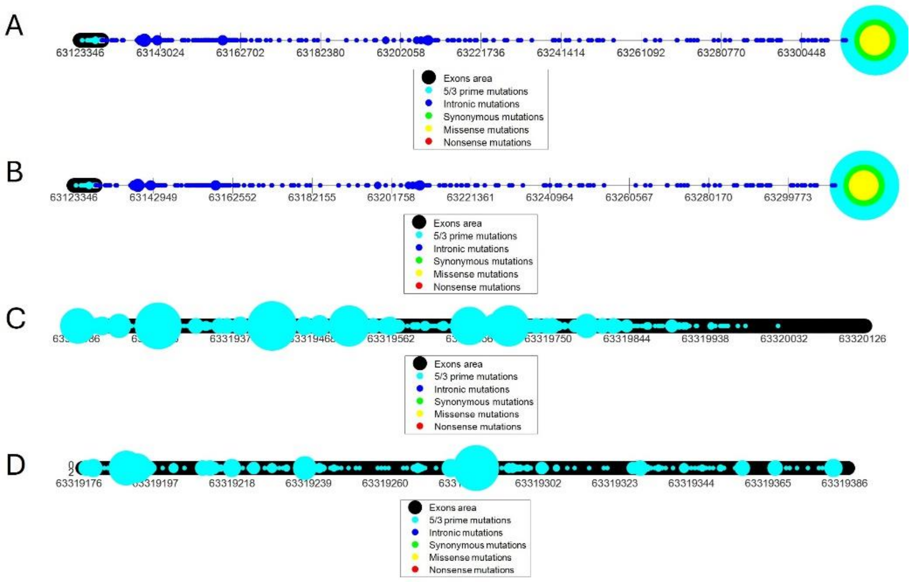


**Fig 17.** This figure illustrates where mutations of DLBCL patients can be found along the BCL2 gene while the size of each circle represents the frequency of the mutation in patients. The figure is divided into four parts, where A shows the mutations along the entire gene. B shows the mutations from the beginning of the gene up until the mutation block in 63319276- 63319286. C shows the mutations from the mutations block in 63319276- 63319286 up until the end of the gene. D shows the mutations in the area of the mutation block (+- 100 nucleotides). As can be seen the 63319276- 63319286 mutations block includes 5' prime mutations where one of them is highly repetitive.

To further validate that this mutation has an effect on the gene, the expression was checked. The expression of the BCL2 gene in patients with the mutation was higher than in randomized groups of malignant lymphoma patients (Pvalue < 0.004 see Fig. 18). We hypothesize that this marked overexpression of BCL2 will be very beneficial for cancer as this is an anti-apoptotic gene.

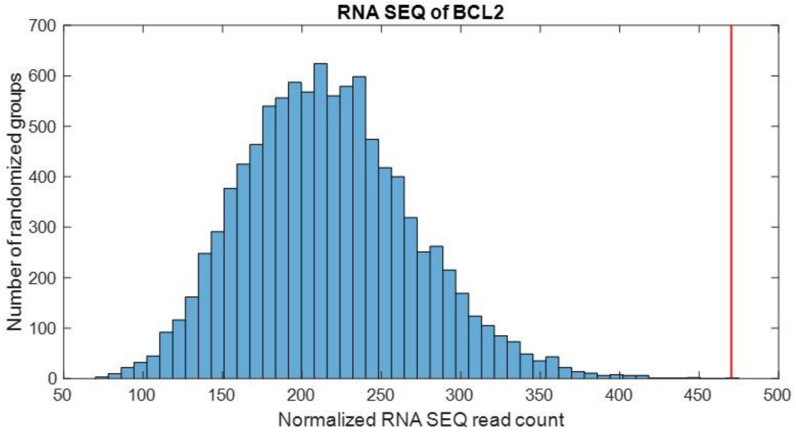


**Fig 18.** Using the RNA sequencing data, 10,000 patient groups were randomly selected, and their mean expression was calculated. The red line represents the mean expression of patients with mutations in the 63319276- 63319286 block in BCL2 gene.

Now a search for relevant transcription factors that might explain the change in the expression was made. Only transcription factors that are expressed in EBV transformed lymphocytes were considered and were checked for a change in the binding ability of the transcription factor to the relevant area. Only one transcription factor was found as a possible cause to the mentioned-above overexpression – BCL2. The mutations in the mutations block interfere with the binding of the MSC repressor to BCL2, as shown in Fig. 19, and by that may cause the overexpression of BCL2.

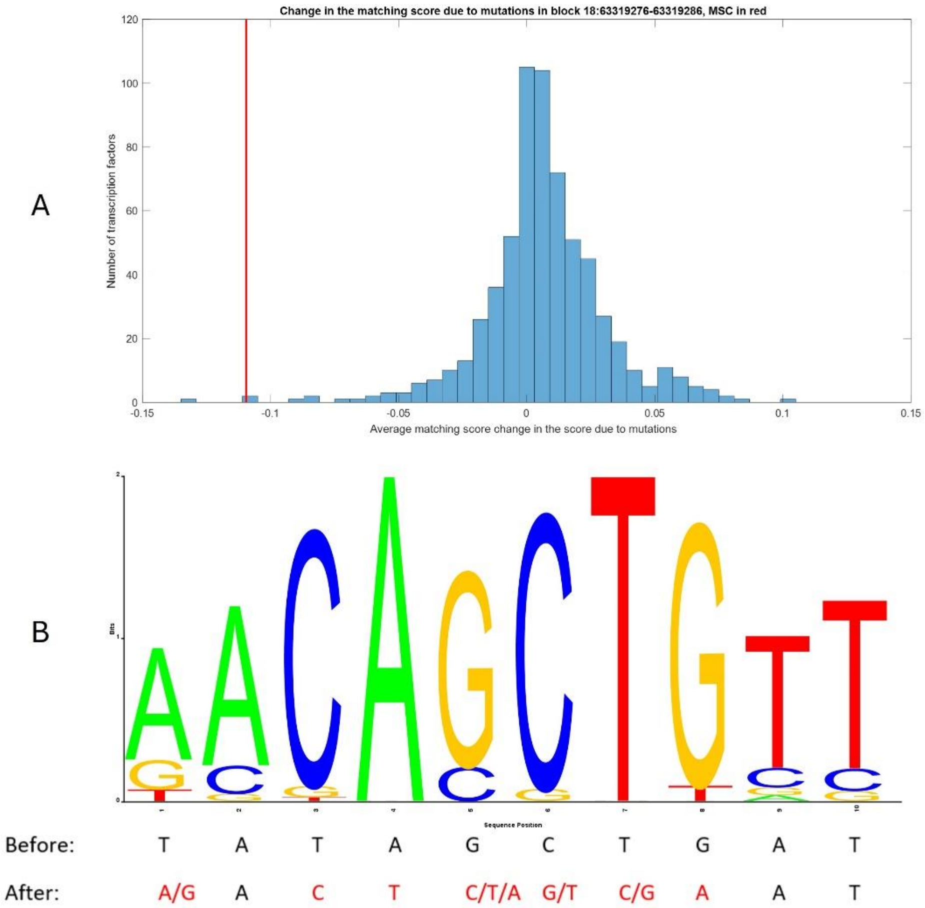


**Fig 19.** Represents the effect the mutations have on the ability of transcription factors to bind to the gene. In A, a histogram representing the influence of the mutation in the block on the ability of each transcription factor to bind. As can be seen, most transcription factors are lowly affected by this mutation, while only a few are highly affected. MSC is marked in red line. In B, The LOGO of the binding site of MSC according to JASPAR's PSSMs. Below the nucleotides of the relevant site (18: 63319278- 63319287 forward strand) is shown with the mutation (after) and without the mutation (before). Marked in red are alternatives nucleotides that were found in patients.

#### References

1. Sang, Y., Kong, P., Zhang, S., Zhang, L., Cao, Y., Duan, X., Sun, T., Tao, Z., Liu, W.: SGK1 in Human Cancer: Emerging Roles and Mechanisms. Front. Oncol. 10, (2021). https://doi.org/10.3389/fonc.2020.608722.
2. Gao, J., Sidiropoulou, E., Walker, I., Krupka, J.A., Mizielinski, K., Usheva, Z., Samarajiwa, S.A., Hodson, D.J.: SGK1 mutations in DLBCL generate hyperstable protein neoisoforms

that promote AKT independence. Blood. 138, 959–964 (2021). https://doi.org/10.1182/blood.2020010432.
3. Augustyn, A., Borromeo, M., Wang, T., Fujimoto, J., Shao, C., Dospoy, P.D., Lee, V., Tan, C., Sullivan, J.P., Larsen, J.E., Girard, L., Behrens, C., Wistuba, I.I., Xie, Y., Cobb, M.H., Gazdar, A.F., Johnson, J.E., Minna, J.D.: ASCL1 is a lineage oncogene providing therapeutic targets for high-grade neuroendocrine lung cancers. Proc. Natl. Acad. Sci. 111, 14788–14793 (2014). https://doi.org/10.1073/pnas.1410419111.
4. Malli, T., Rammer, M., Haslinger, S., Burghofer, J., Burgstaller, S., Boesmueller, H.-C., Marschon, R., Kranewitter, W., Erdel, M., Deutschbauer, S., Webersinke, G.: Overexpression of the proneural transcription factor ASCL1 in chronic lymphocytic leukemia with a t(12;14)(q23.2;q32.3). Mol. Cytogenet. 11, 3 (2018). https://doi.org/10.1186/s13039-018-0355-7.
5. Slack, G.W., Gascoyne, R.D.: MYC and Aggressive B-cell Lymphomas. Adv. Anat. Pathol. 18, (2011).
6. Aguilar-Medina, M., Avendaño-Félix, M., Lizárraga-Verdugo, E., Bermúdez, M., Romero-Quintana, J.G., Ramos-Payan, R., Ruíz-García, E., López-Camarillo, C.: SOX9 Stem-Cell Factor: Clinical and Functional Relevance in Cancer. J. Oncol. 2019, 6754040 (2019). https://doi.org/10.1155/2019/6754040.
7. Shen, Y., Zhou, J., Nie, K., Cheng, S., Chen, Z., Wang, W., Wei, W., Jiang, D., Peng, Z., Ren, Y., Zhang, Y., Fan, Q., Richards, K.L., Qi, Y., Cheng, J., Tam, W., Ma, J.: Oncogenic role of the SOX9-DHCR24-cholesterol biosynthesis axis in IGH-BCL2+ diffuse large B-cell lymphomas. Blood. 139, 73–86 (2022). https://doi.org/10.1182/blood.2021012327.
8. Gordon, S., Akopyan, G., Garban, H., Bonavida, B.: Transcription factor YY1: structure, function, and therapeutic implications in cancer biology. Oncogene. 25, 1125–1142 (2006). https://doi.org/10.1038/sj.onc.1209080.
9. Castellano, G., Torrisi, E., Ligresti, G., Nicoletti, F., Malaponte, G., Travali, S., McCubrey, J.A., Canevari, S., Libra, M.: Yin Yang 1 overexpression in diffuse large B-cell lymphoma is associated with B-cell transformation and tumor progression. Cell Cycle. 9, 557–563 (2010). https://doi.org/10.4161/cc.9.3.10554.
10. Shami-Schnitzer, O., Zafir, Z., Tuller, T.: Novel Driver Synonymous Mutations in the Coding Regions of GCB Lymphoma Patients Improve the Transcription Levels of BCL2. In: Bebis, G., Alekseyev, M., Cho, H., Gevertz, J., and Rodriguez Martinez, M. (eds.) Mathematical and Computational Oncology. pp. 108–118. Springer International Publishing, Cham (2020).
11. Cory, S., Adams, J.M.: The Bcl2 family: regulators of the cellular life-or-death switch. Nat. Rev. Cancer. 2, 647 (2002).
12. Monni, O., Franssila, K., Joensuu, H., Knuutila, S.: BCL2 Overexpression in Diffuse Large B-Cell Lymphoma. Leuk. Lymphoma. 34, 45–52 (1999). https://doi.org/10.3109/10428199909083379.
